# Dissecting the neuronal mechanisms of invariant word recognition

**DOI:** 10.1101/2023.11.06.565812

**Authors:** Aakash Agrawal, Stanislas Dehaene

## Abstract

Learning to read places a strong challenge on the visual system. Years of expertise lead to a remarkable capacity to separate highly similar letters and encode their relative positions, thus distinguishing words such as FORM and FROM invariantly over a large range of sizes and absolute positions. How neural circuits achieve invariant word recognition remains unknown. Here we address this issue through computational modeling and brain imaging. We first trained deep neural network models to recognize written words, then analyzed the reading-specialized units that emerged in deep layers. With literacy, units became sensitive to specific letter identities and their distance from the blank space at the left or right of a word, thus acting as “space bigrams” encoding ordinal position using an approximate number code. Using 7T functional MRI and magnetoencephalography in adults, we localized the predicted ordinal code anatomically (visual word form area) and temporally (∼220ms). The proposed neuronal mechanism for invariant word recognition can explain reading errors and makes precise predictions about how position-invariant neural codes arise in brains and artificial systems.

## Introduction

Distinguishing two similar visual shapes, and doing so regardless of huge variations in superficial appearance, is a problem that the human ventral visual pathway solves with remarkable efficiency. This capacity for invariant visual recognition is particularly evident in the case of fluent reading. 3 or 4 times per second, a fluent reader lands eyes on a word and reliably recognizes it among tens of thousands of similar entries in the mental lexicon. A specific challenge raised by fluent reading is the need to distinguish words such as FROM and FORM that only differ in relative letter position (Grainger & Whitney, 2004) – and to do so invariantly over a wide range of fonts, sizes, spacings, and retinal positions (Legge & Bigelow, 2011; Vinckier et al., 2011; Xiong et al., 2019). Here, we propose and test a hypothesis about the neuronal circuit that solves this problem.

In recent years, the cortical areas underlying fluent reading have begun to be resolved. The acquisition of literacy leads to the formation of a specialized word-responsive region in the ventral visual cortex, the Visual Word Form Area (VWFA) (Dehaene et al., 2010, 2015; Dehaene-Lambertz et al., 2018). Brain imaging shows that this region becomes attuned to stimuli in the learned script (C. I. Baker et al., 2007; Szwed et al., 2011, 2014) and preferably responds to stimuli that respect the distributional statistics of letters in the learned language (Binder et al., 2006; Vinckier et al., 2007; Woolnough et al., 2020; Zhan et al., 2023). The VWFA responds in a largely invariant manner to identical words that vary in case and location across the visual field (Cohen et al., 2002; Dehaene et al., 2001, 2004), although it remains weakly modulated by absolute position (Rauschecker et al., 2012). Simultaneously, the VWFA differentiates anagrams such as RANGE and ANGER that differ only in the order of their letters (Dehaene et al., 2004).

Despite those advances, the nature of the neural code underlying this invariant recognition remains unknown. Two broad classes of theories can be opposed (McCloskey et al., 2013). Contextual schemes assume that letters are encoded relative to the location of other letters. For instance, the visual system may extract bigrams, i.e., ordered pairs of letters. Thus, FORM and FROM would be distinguished by their bigrams “OR” versus “RO”, regardless of where they occur on the retina. Encoding the most frequent such bigrams would suffice to recognize many words (Dehaene et al., 2005; Grainger & Whitney, 2004; Whitney, 2001). Positional schemes, on the other hand, assume that words are encoded by a list of letters, each attached to its relative ordinal location within the word (Coltheart et al., 2001; McClelland & Rumelhart, 1981; Norris, 2013). For instance, each letter may bear a neural code for its approximate ordinal number relative to the beginning of the word (see McCloskey et al., 2013 for a list of relative positional schemes).

Functional MRI data initially supported the bigram hypothesis by revealing stronger VWFA responses to stimuli containing frequent letter bigrams (Binder et al., 2006; Vinckier et al., 2007). However, recently, evidence from psychophysics and time-resolved intracranial recordings suggests that frequent bigrams may not contribute to recognition, at least during the first ∼300 ms of word recognition, where only frequent letters are separated from rare letters or non-letters (Agrawal et al., 2020; McCloskey et al., 2013; Woolnough et al., 2020). What develops with literacy is the compositionality of visual word representation, which increases the dissimilarity and independence between individual letters at nearby locations (Agrawal et al., 2019, 2022). Compared to contextual bigram coding, ordinal letter coding provides a better fit to the psychophysical distance between letter strings (Agrawal et al., 2020), the early intracranial responses to written words (Woolnough et al., 2020), and the responses of word selective units in artificial deep networks (Hannagan et al., 2021).

What remains to be understood is how such an invariant ordinal code is achieved by neural circuits. Most cognitive models of reading simply beg the question by taking as input a bank of position-specific letter detectors, responding for instance to letter R in 2^nd^ ordinal position (Coltheart et al., 2001; McClelland & Rumelhart, 1981; Norris, 2013), and implicitly assuming that some earlier unknown mechanism normalized the input for size, font, and retinal position to eventually encode letters in a relative (ordinal) rather than absolute (retinal) spatial reference frame.

Resolving the neural code for reading is difficult, due to the poor resolution of functional magnetic resonance imaging (fMRI) and the lack of animal models for detailed neurophysiological investigations (although see (Grainger et al., 2012; Rajalingham et al., 2020; Ziegler et al., 2013)). Here, we show that precise predictions about the neurophysiological architecture for invariant reading can be obtained by studying convolutional neural networks (CNN) models of the ventral visual cortex that, like literate humans, can be recycled for reading. We trained CNNs to recognize written words in different languages and analyzed their responses to both trained and novel scripts. After validating those models against earlier psychophysical studies (Agrawal et al., 2019), we investigated how they achieve invariant word recognition by characterizing their units’ receptive fields to letters and strings at various stages. This led us to discover a novel principle of relative position coding, “space bigrams”. Finally, we validated the predictions of the proposed model and mapped its stages of processing in the human brain using high-resolution 7T fMRI and magnetoencephalography (MEG).

## Results

### Invariant word identification

We first trained various instances of CORnet-Z, a CNN whose architecture partially matches the primate ventral visual system (Kubilius et al., 2018), to recognize 1000 images (illiterate networks), then an additional 1000 words in different writing systems (literate networks; Figure 1A). The networks trained on ImageNet reached accuracy levels that were comparable to the earlier reported values for CORnet-Z (top-1 accuracy = 36.8% ± 0.1). With the introduction of words, performance on ImageNet dropped marginally (top-1 accuracy = 36.4% ± 0.4) while becoming excellent on word recognition across different languages, with test words varying in case, font, location, and size (see methods) (top-1 accuracy = 88.2% ± 0.5 for French, 87.5% ± 0.5 for English, 92.2% ± 0.3 for Chinese, 95.5% ± 0.2 for Telugu, 90.9% ± 0.5 for Malayalam). Networks trained on bilingual stimuli reached accuracy levels comparable to monolingual networks (top-1 accuracy = 88.6% ± 0.7 for the English + French network, and 91.0% ± 0.2 for the English + Chinese network). Higher accuracy with words than with images can be attributed to the limited variations of text on a plain background.

**Figure 1:**
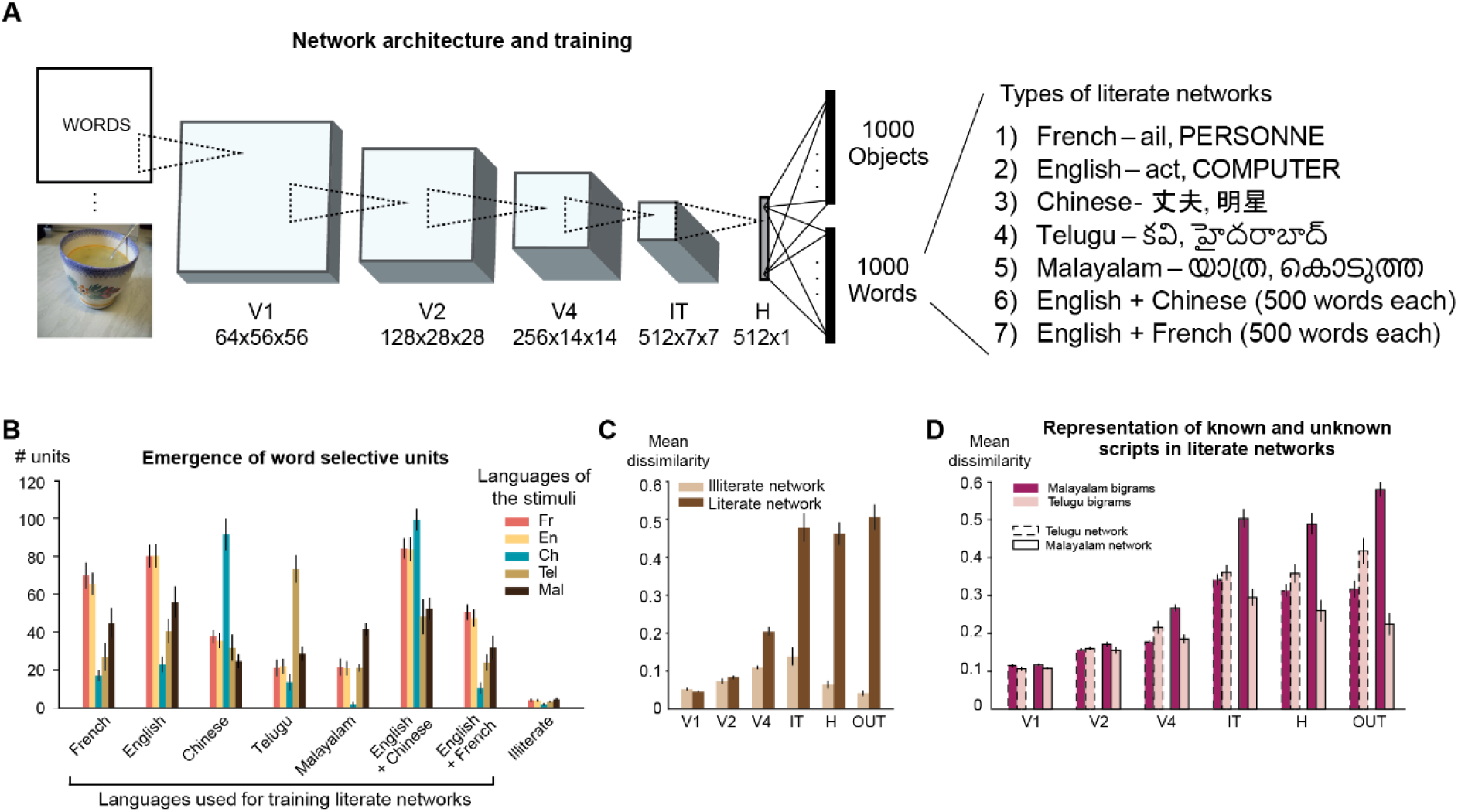
Properties of a neural network trained to recognize words. (A) The CORnet-Z architecture was used as a model of the ventral visual pathway. The illiterate network was trained to predict 1000 object categories present in the ImageNet dataset. The literate networks were trained to predict both the 1000 object categories and 1000 word categories from a given language. (B) The number of word-selective units in the H layer of different literate and illiterate networks for both trained and novel scripts. Error bars indicate standard deviation across 5 different instances. (C) Mean dissimilarity estimated across bigrams pairs (n = ^49^C_2_ = 1176) from different layers of French literate (dark) and illiterate (light) networks (similar results were obtained in networks trained with other languages). (D) Mean dissimilarity estimated across Telugu (n = ^25^C_2_ = 300) and Malayalam (n = ^25^C_2_) bigram pairs from different layers of Telugu (dashed border) and Malayalam (solid border) networks

### Emergence of script-specific units

We next tested for the emergence of units specialized for the visual form of words, similar to the human VWFA. Within the non-convolutional, penultimate layer of each network (h), we searched for units selective to a given script over and above other categories such as faces or objects, a contrast similar to fMRI studies of the VWFA (see methods). Literacy dramatically enhanced the number of script-selective units, from a mean of 4.2 units in illiterate networks to 40-100 units (Figure 1B). Relative to our previous work (Hannagan et al., 2021), where a single script was probed, here we could evaluate the selectivity to the trained script relative to others. In agreement with human VWFA fMRI (C. I. Baker et al., 2007), many more units responded to the trained script than to untrained ones (Figure 1B). Nevertheless, in literate relative to the illiterate networks, greater responses were also seen to untrained scripts. Although few, there were units in each network that exhibited selectivity to novel scripts but not the trained script (n = 3 for French network, 10 for English, 6 for Chinese, 3 for Telugu, 5 for Malayalam, 17 for English + Chinese, 5 for English + French). This finding mimics the experimental observation that, while preferring the learned script(s), the VWFA also responds at a lower level to unknown scripts (C. I. Baker et al., 2007; Szwed et al., 2011, 2014).

Among the units that were selective to the trained script, their response to unknown scripts depended on their similarity to the trained script. For instance, the English literate network has a greater number of Malayalam selective units compared to Chinese selective units, presumably due to the presence of rounded symbols in both scripts (Figure 1B). Furthermore, in “bilingual” networks trained with two scripts, consistent with fMRI of bilingual readers (Zhan et al., 2023), a large proportion of word-selective units were common to the two trained languages, yet more so when the two languages shared the same alphabet (English-French network: 99.8% of French word selective units, and 100 % of English units, also responded to the other script) than when they did not (English-Chinese network: only 75% of Chinese word selective units responded to English, and vice-versa for 84% of English units).

### Improvements in neural discriminability

Literacy leads to script-specific behavioral and neural response enhancements at both early and late visual stages (Chang et al., 2015; Dehaene et al., 2010, 2015; Szwed et al., 2012) and, in particular, increases the perceived dissimilarity between letters, bigrams and words (Agrawal et al., 2019). To examine whether this effect was present in our simulations, we compared the mean pair-wise neural dissimilarity between 49 bigrams in French literate and illiterate networks (Figure 1C; see methods). From the V4 layer on, the mean dissimilarity was indeed higher for the literate than for the illiterate network, indicating that the neural population had improved its representation of letter combinations. We also repeated the 2x2 analysis reported by (Agrawal et al., 2019), which compared the mean dissimilarity between Telugu and Malayalam words in readers of either script. Our simulations replicated the finding that visual dissimilarity is higher for the learned script, and localized this effect to mid-visual areas, starting in area V4 (Figure 1D). Similar effects were found with networks trained in other languages; to avoid repetition, for the remnant of the paper, we focused on networks trained to recognize French words (the language in which we scan participants in fMRI and MEG).

### Letter tuning

Using 1000 words with variable length, we replicated our previous observation that a large amount of variance in the activity of word-selective units in the penultimate H layer could be captured by a letter X position encoding model (Hannagan et al., 2021). Some units care about a single letter regardless of its position, while most units care about one or several letters at a specific ordinal position (Figure 2A). A model-based comparison of various letter-position schemes revealed improved fits when units were assumed to fire at a fixed ordinal position relative to word beginning or ending, rather than relative to either alone or to word center (Figure 2B), in agreement with human behavior (McCloskey et al., 2013). Position tuning was sharper near those edge letter positions (i.e., 1^st^, 2^nd^, penultimate, or end position) compared to middle letter positions (Figure 2C). These findings align with actual recordings of number neurons (Nieder & Dehaene, 2009) and with previous behavioral studies that found a greater sensitivity of human readers to detect changes in edge letters than in middle ones (Agrawal et al., 2020; Grainger & Whitney, 2004).

**Figure 2:**
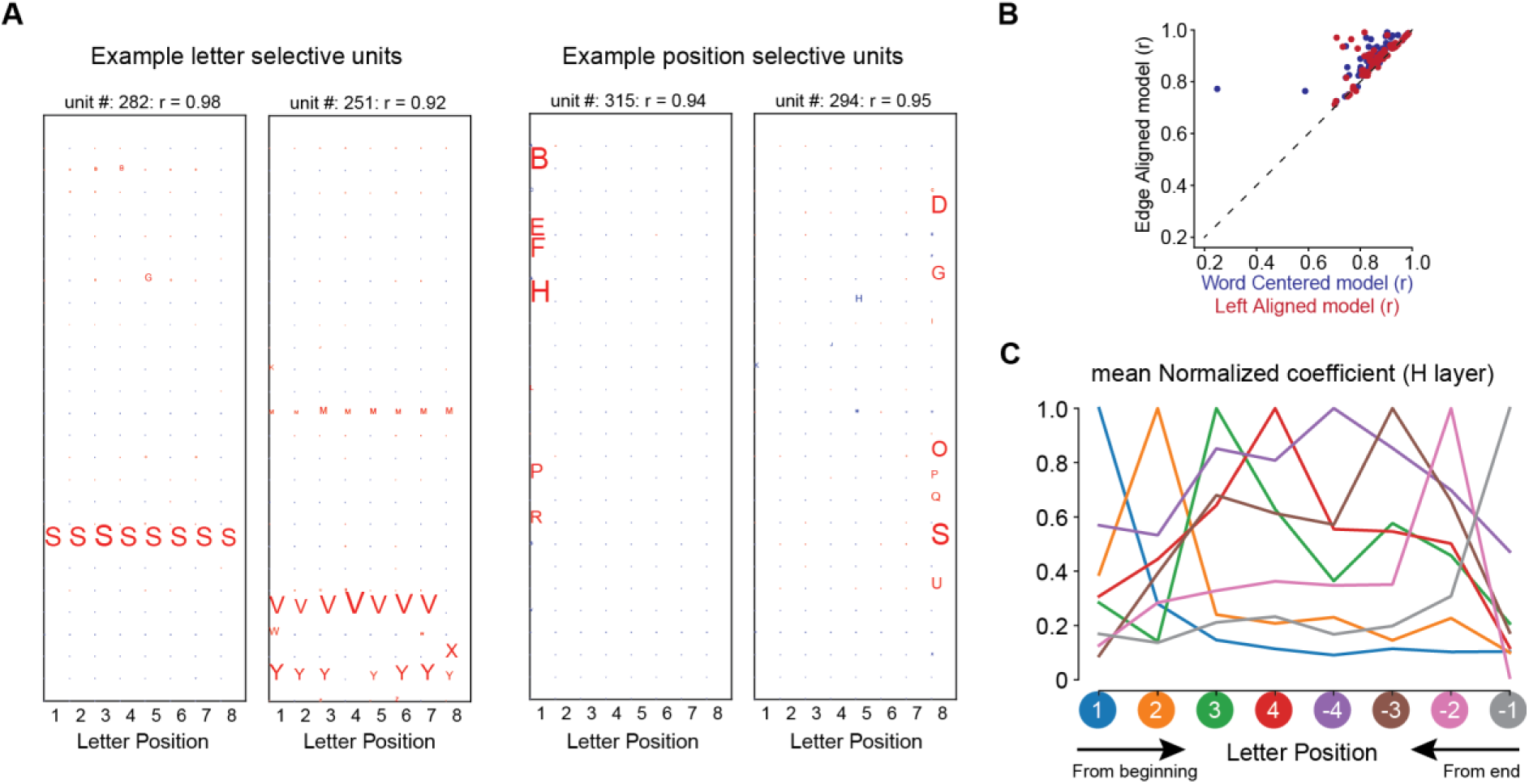
Position encoding schemes. (A) Visualization of the letter model coefficients for a few example units. The size of the letter indicates the magnitude. (B) Comparison between the model fits across all word selective units in the H layer using Edge aligned position coding scheme and word-centered (blue) & left-aligned model fits (red). The dashed line represents the unity slope line. (C) Average coefficients of the ordinal-position regression model in the H layer. Following (Nieder & Dehaene, 2009), units were sorted according to their preferred ordinal location from word beginning or ending (colors), and the coefficients of each unit were normalized by dividing by their maximum value and then averaging across units.

### Mechanisms of invariant word recognition

How did units in the last layer acquire such invariant ordinal tuning, from a convolutional network whose initial layers have absolute (retinotopic) receptive fields? To dissect the reading circuit, we first identified, at each layer of the French literate network, the units preferring words over other stimuli such as faces or objects (see methods). The proportion of word selective units increased with successive layers (percentage of units = 0.02% in V1, 0.6% in V2, 1.76% in V4, 3.61% in IT, 13.6% in h). To evaluate the units’ receptive fields, we next tested each letter-selective unit with stimuli designed to dissociate retinotopic letter position, word position, and ordinal letter position codes (Figure 3). First, for each unit, we identified its most and least preferred letter (see methods). These letters were then used to create 4-letter stimuli where a single preferred letter (e.g., o) was embedded at various locations within a string of non-preferred letters (e.g., xoxx). In our 5x4 factorial design, retinotopic word position (5 levels) varied across the rows, while ordinal position of the preferred letter (4 levels) varied across the columns (Figure 3a). Across that 5x4 stimulus matrix, units coding for a fixed retinotopic letter position should exhibit a diagonal response profile (Figure 3B-left). Conversely, units that encode a fixed ordinal position, regardless of word position, will have a vertical response profile (Figure 3B-right).

**Figure 3:**
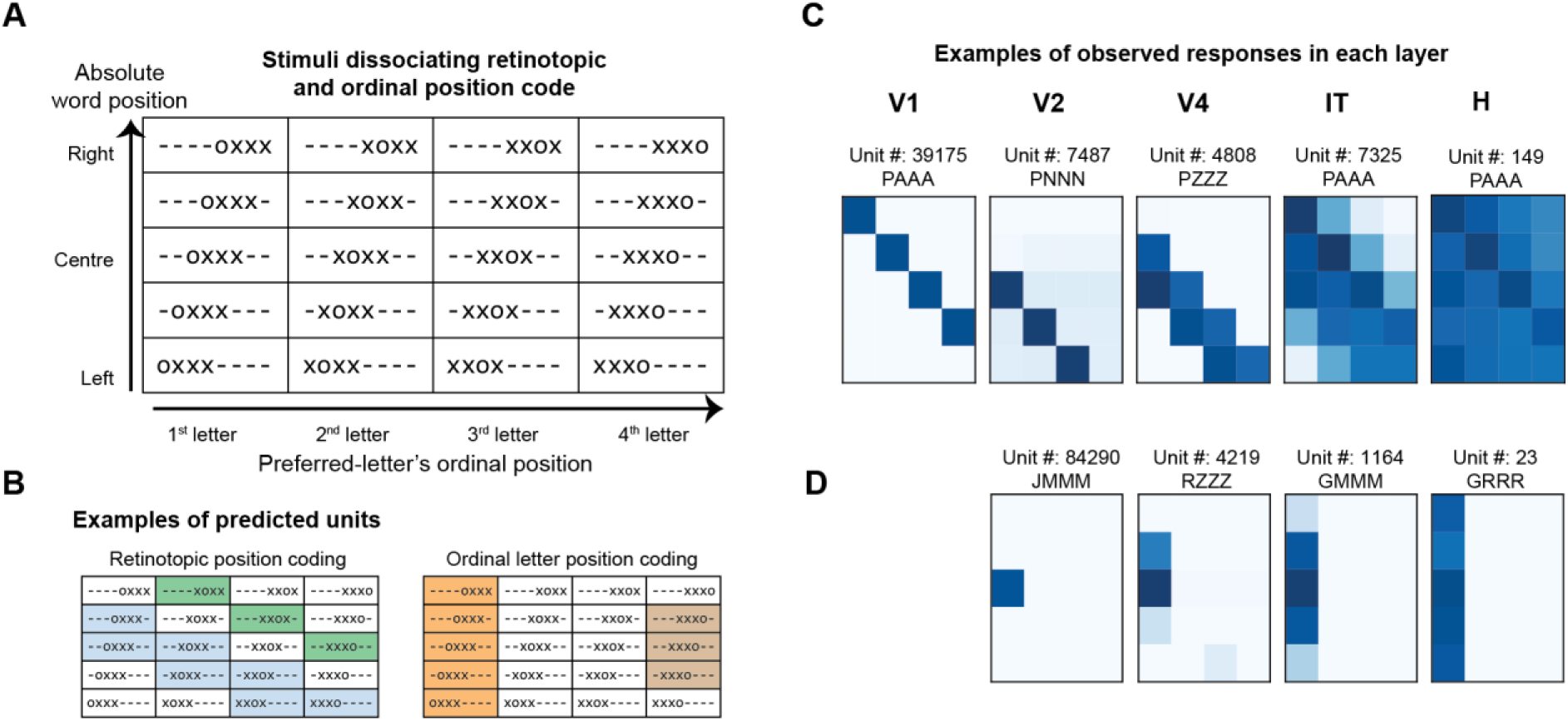
Transition from absolute (retinotopic) coding to ordinal position coding. (A) Schematic of the stimuli used to dissociate absolute vs relative coding. We presented a single preferred letter within a “word” made of multiple non-preferred letters. Here, the preferred letter is ‘o’, and the unpreferred letter is ‘x’. Absolute word position varies across rows, while ordinal preferred-letter position within a string varies across columns. ‘-’ represents blank space. (B) Expected response profile for two types of units with (1) absolute, retinotopic letter position coding (left); and (2) relative, ordinal letter-position coding (right). Each color represents a different unit. (C) Exemplar units from each layer of the French literate network. The unit-id and the string comprising preferred and unpreferred letter is displayed on the top. Darker shades represent a higher response. (D) Same as (C) but for units showing ordinal position coding.

In our networks, we indeed observe units that were sharply tuned to either retinotopic or ordinal positions. The proportion of retinotopic units was highest in the initial layers of the network (n = 100% in V1, 46.5% in V2, 56% in V4, 20.8% in IT, and 3.4% in H layer). Conversely, the later layers of the network formed units that encoded the ordinal position of the preferred letter (83.1% in the H layer).

This analysis uncovered two main types of units (examples from each layer are shown in Figure 3C-D). In both cases, as expected, receptive field size increased along the successive layers of the network, and units in the V1 layer responded to the preferred letter only when presented at a specific position. However, the first type of units continued to respond to a single letter over a progressively increasing range of retinotopic positions, with mid-layer units spanning 2-4 letter positions, and the h units showed complete position invariance (Figure 3C). These units are easy to understand: by pulling over progressively larger receptive fields, each layer achieves greater position invariance in recognizing a specific letter. However, such units are unable to encode letter order and separate anagrams. More interestingly, other units in the intermediate layers of the network responded only when the preferred letter was presented within the given receptive field *and* at a fixed ordinal position, often either the first or the last position in a word (n = 7.8 % in V2, 11.8 % in V4, 37.3 % in IT). Such units are referred as “edge coding units” and were observed in all layers but V1 (Figure 3D). Across V2, V4, and IT, these units exhibited a broadening of their retinotopic receptive fields, until a complete invariance to retinotopic position and a pure selectivity to ordinal position were attained in the h layer (Figure 3D). Thus, ordinal coding was achieved by pooling over a hierarchy of edge-sensitive letter detectors. While this scheme was dominant, we also observed units with mixed selectivity that responded to a diverse range of positions that were neither purely retinotopic nor ordinal (Figure S4).

Next, we investigated the origins of those crucial edge coding units. We hypothesized that, within the convolutional layers, units managed to encode approximate ordinal position because their convolution field was jointly sensitive to (1) one or several specific letter shapes, *and* (2) the presence of a blank space (absence of any letter), either to the left (for units coding ordinal position relative to word beginning) or to its right (for end-coding units). We term these units “space bigrams” because they are sensitive to a pair of characters, one of which is a space. The space bigram coding scheme is therefore an extension of the previous open-bigram hypothesis, which assumed that units would be sensitive to ordered letter pairs such as “OR”, by allowing one of these letters to be a space (and space is indeed the most frequent character (∼20%) in English, French, and probably all similar alphabetic codes).

To separate space coding from ordinal position coding, we tested our network’s responses to a modified stimulus set where a blank space was introduced between the letters (e.g., x o x). Behavioral and brain-imaging tests show that inserting a single s p a c e between letters does not disrupt normal reading (Cohen et al., 2008; Vinckier et al., 2011) and may even facilitate it (Perea & Gomez, 2012; Zorzi et al., 2012). To maintain the overall count of possible letter positions, the number of letters in a given stimulus was reduced to three, consisting of one preferred and two non-preferred letters. This resulted in a 4x3 stimulus matrix, where absolute word position (4 levels) varied across rows, and the ordinal position of the preferred letter varied across the columns (3 levels; Figure 4A). If edge-coding units were indeed encoding the presence of a nearby blank space rather than a genuine ordinal position, their responses would persist even when the preferred letter was present at the middle location, provided it was flanked by a blank space (Figure 4B-top). Conversely, a genuine ordinal position unit would continue to respond to the same ordinal location even in the spaced condition (Figure 4B-bottom).

**Figure 4:**
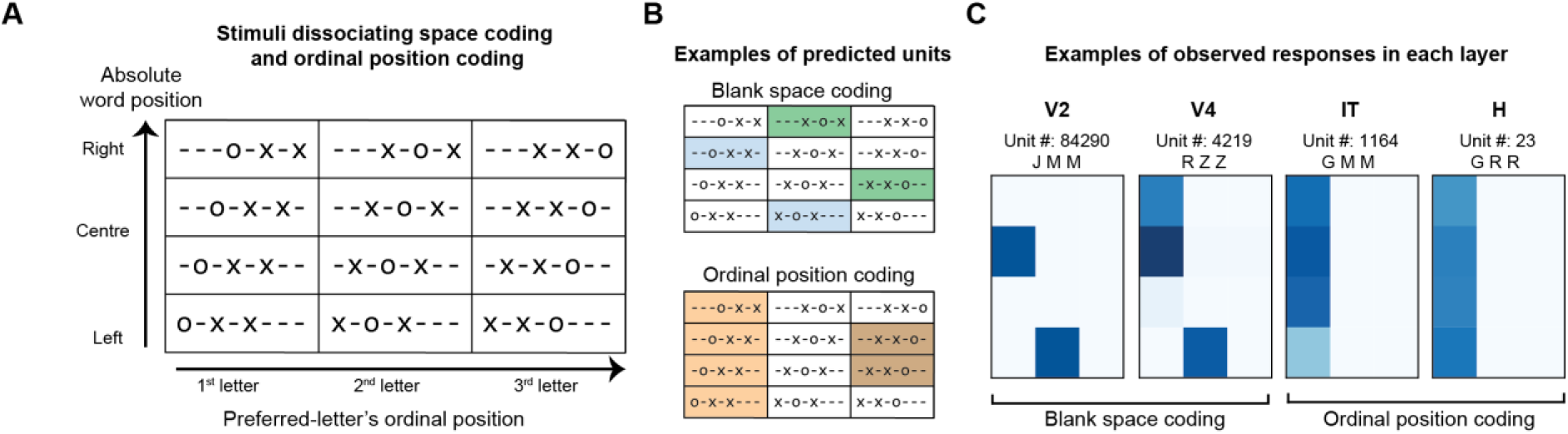
Transition from blank space to ordinal position coding. (A) Same as Figure 3 but with stimuli used to dissociate blank space vs ordinal position coding units. (B) Expected response profile of units with blank space coding (top) and ordinal letter position coding (bottom). Each color represents a different unit. (C) Exemplar units from each layer of the French literate network. The unit-id and the string comprising preferred and unpreferred letter is displayed on the top. Darker shades represent a higher response.

Figure 4C shows the responses of the same units as in Figure 3D when tested using the spaced stimulus set. We found a clear transition across layers: V2 and V4 units coded for blank spaces, while IT and H layers encoded ordinal positions by pulling over several such blank space units. We could explain mechanistically how those units work, based on their connectivity patterns (see Figure S5 and corresponding supplementary text). For instance, a specific unit (# 84290) in the V2 layer exhibited maximal response when its preferred letter ‘J’ was presented at the beginning of the word, with a receptive field centered on the third retinotopic letter position (Figure 3D). When tested with spaced stimuli, the unit maintained its receptive field location (i.e., third retinotopic position) and continued to respond to the letter J, but now did so whenever it was preceded by a blank space, even if it was not located at the beginning of the word. Units in the V4 layer exhibited a comparable response pattern. However, in the IT layer, units maintained their preference for the first ordinal position even within s p a c e d strings. These findings provide robust evidence that edge coding units initially encode blank spaces and contribute to the ultimate extraction of ordinal position coding.

### Optimal stimuli for each layer

To study the properties of neural networks, a complementary approach consists of identifying the most and the least preferred stimuli for individual units. This investigation can be performed using a technique known as neural network distillation, which is based on image-level gradient descent. By iteratively adjusting the pixel values of an image, a stimulus is generated that maximizes/minimizes the activation of a specific unit. Analyzing these generated images enables us to identify the distinguishing features preferred by each unit.

Here, we started with 3 words of varying length (AIR, PAIN, and SQUARE) and found the unit with the highest activation in each layer. Next, for each unit, we generated its most preferred stimuli using the network distillation toolbox (see methods). In the early layers (V1 and V2), units preferred dark-oriented lines and curved shapes against a white background (Figure 5), presumably reflecting the background of the training word dataset. Consistent with earlier reports (Zeiler & Fergus, 2013), feature complexity and receptive field size increased in later layers of the network. Remarkably, from IT and output, the automated image optimization process recovered word fragments that partially matched the word identity originally used to select the units (Figure 5). Furthermore, the optimal stimuli repeated those fragments over the visual scene, with an inter-stimulus spacing of at least one letter, thus reflecting the eventually complete spatial invariance of the output units.

**Figure 5:**
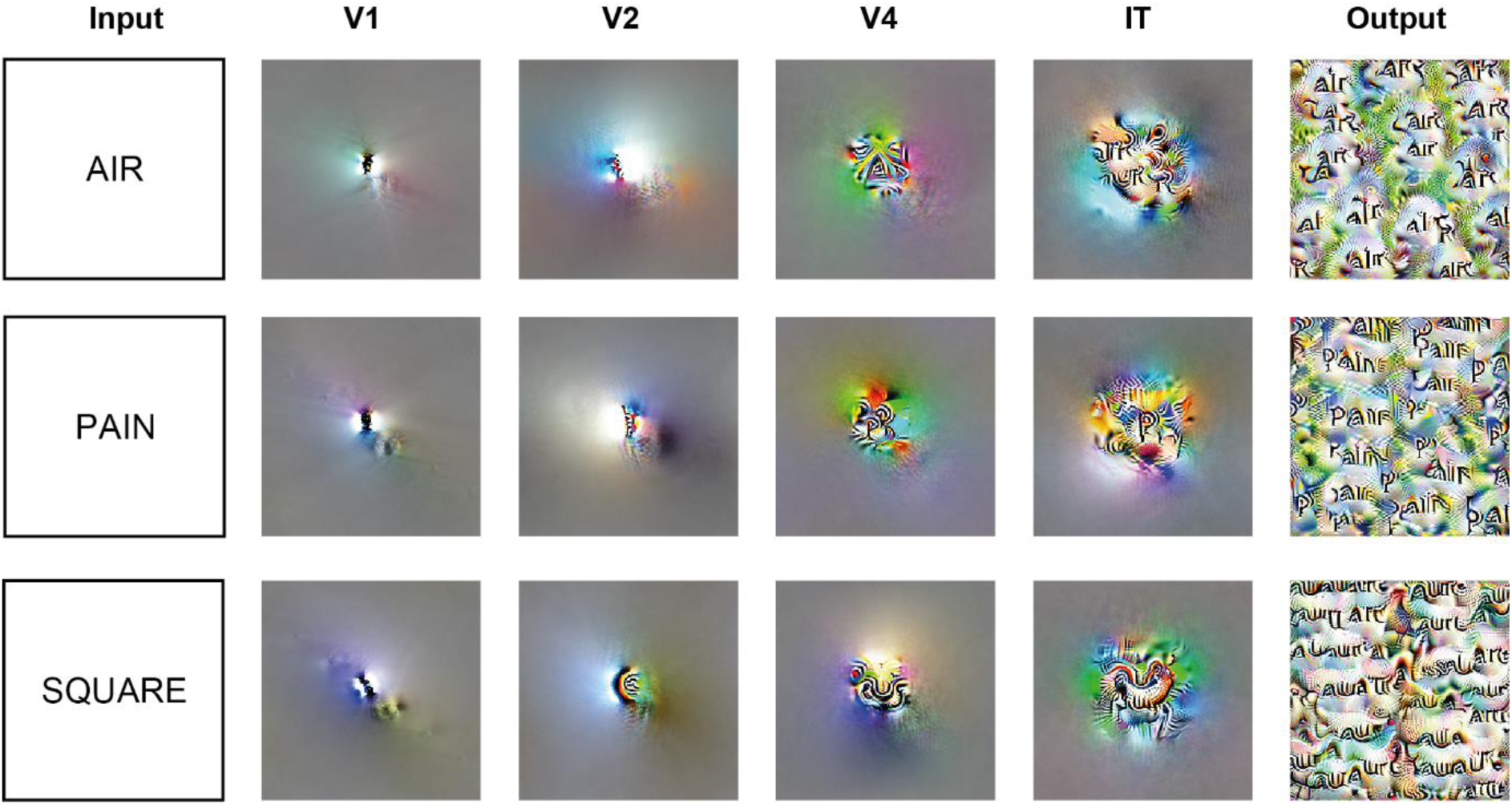
Activation-maximization of word selective units. For each word (input), the features of the channel whose units evoked the highest response within a given layer are shown. For visualization purposes, features are displayed at the central location. Since the H layer is formed by the average pooling of IT layer units, its features are identical to the IT layer and are therefore excluded.

### Letter position coding in the human brain

We next examined whether the progression from retinotopic to ordinal coding predicted by our model could be observed in the brains of expert readers. We recorded neural signals using fMRI and MEG while subjects viewed stimuli similar to those used to investigate literate networks (Figure 3; see methods). Unlike neural network models, however, it is not feasible to map the preferred letter(s) for each voxel in the brain. Therefore, we restricted stimulus selection to three letter pairs comprising a frequent letter and another rare dissimilar one in French (OX, TB, and EQ). We further restricted the stimulus length to three-letter pseudowords. We therefore collected activation patterns for a total of 36 stimuli (3 letter pairs X 3 ordinal positions of the frequent letter X four retinotopic word positions; figure S1).

The stimuli were presented in random order, with the instruction to fixate a red dot centered on the screen, and to press a button on trials (10%) in which all letters were identical (XXX, BBB, or QQQ). Participants performed this task accurately (Accuracy: 94.2% in fMRI, and 81.1% in MEG), and were faster and more accurate when the test stimulus was closer to the fovea (figure S2).

From whole-brain 7T fMRI data (n = 16; 1.2 mm isotropic), using a combination of functional localizer and anatomical maps, we extracted four individual regions of interest (ROIs) along the ventral visual pathway: V1-V3, V4, LOC, and VWFA (see methods). In each region, using voxels as features, we fitted a linear classifier to decode the following factors: word position, letter identity, absolute letter position, and ordinal letter position. To avoid overfitting, we performed 3-fold cross-validation (see methods; MEG data). Consistent with earlier findings, decoding accuracy for word position was highest in early visual areas, but progressively decreased along the ventral visual pathway, ultimately becoming statistically insignificant within VWFA (Figure 6). Contrarily, the decoding of ordinal position was significant only within VWFA. The other two factors could be significantly decoded across all ROIs, with higher accuracy for letter identity in object and word selective regions than early visual areas (p < 0.005; paired t-test).

**Figure 6:**
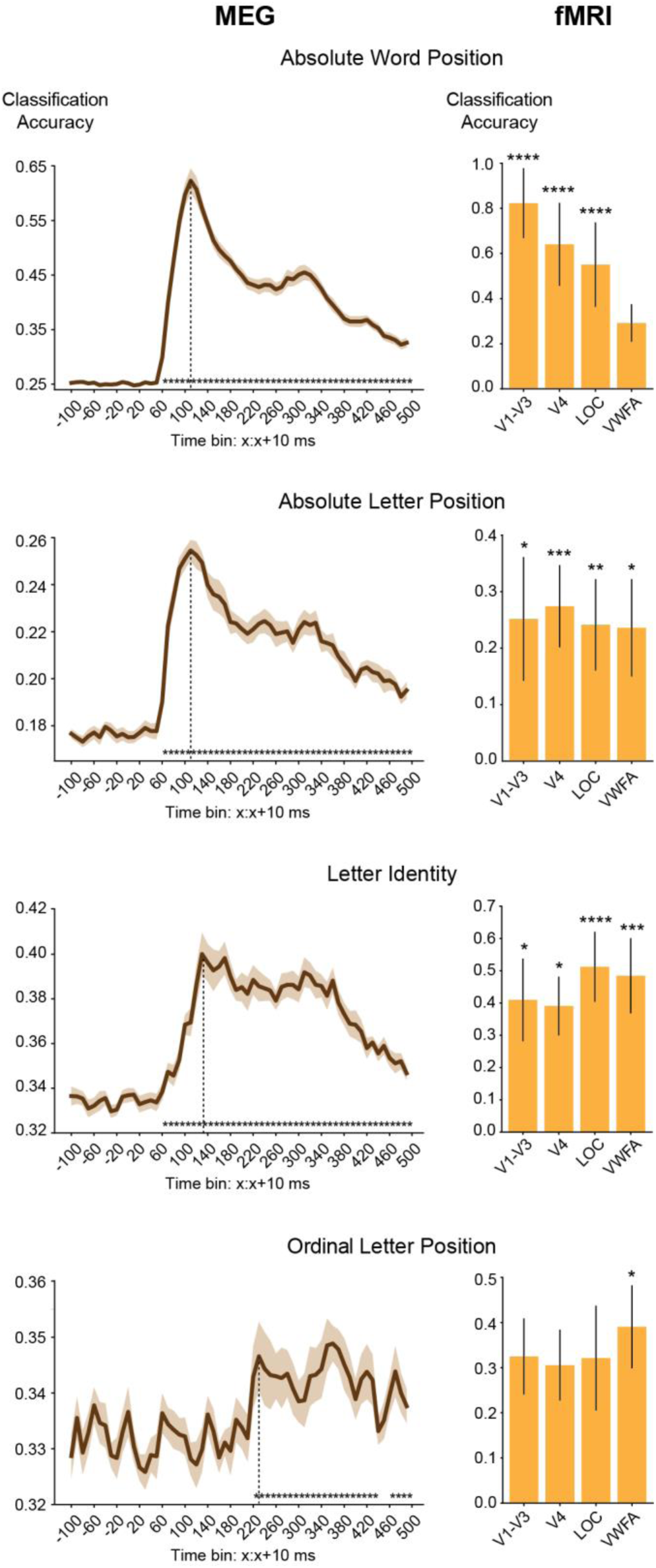
Decoding stimulus properties across space and time. Classification accuracy across different time points from MEG (left) and ROIs from fMRI (right) for absolute word position (chance = 25%), absolute letter position (chance = 16.67%), letter identity (chance = 33.33%), and ordinal letter position (chance = 33.33%). Error bars indicate standard error of mean across subjects and the asterisks indicate statistical significance (in fMRI, * p < 0.05, ** p < 0.005, *** p < 0.0005, etc. using one sample t-test; in MEG, * p < 0.05 using cluster-permutation test).

To further compare the stages of processing between networks and humans, we computed representation dissimilarity matrices (RDM) in each ROI and correlated them with the RDMs estimated from different layers of the literate network. Consistent with earlier studies using objects (Güçlü & Gerven, 2015), the representation in early visual areas best correlated with representation in early layers of the network, and the representation in VWFA best correlated with the later layers of the network, with a peak at IT layer (Figure S9). A whole-brain searchlight analysis further confirmed these observations and revealed that, beyond VWFA, the later layers of the literate network also matched with the representation space of higher areas involved in language processing (Supplementary Section 3).

Next, to investigate the neural dynamics of letter position coding, we recorded 306-channel MEG signals to the same stimuli in a different group of participants (n = 17). MEG signals were grouped into bins of 10ms each, which were submitted to the same decoding and representational similarity analyses. Decoding accuracy was above chance for word position, absolute letter position, and letter identity starting at ∼60ms and peaking around ∼140ms. However, decoding of ordinal letter position became significant only at ∼220ms, which aligns with the timings of word selectivity in the anterior region of the fusiform gyrus (Thesen et al., 2012). Further, the representation space in later layers of CNN was also significantly correlated with the representation of MEG signals around the same time window (Figure S10).

Overall, our findings indicate that while early visual signals are driven by retinotopy, the VWFA encodes ordinal letter position.

## Discussion

Most cognitive models of reading assume a letter X position code as input (Norris, 2013), yet without showing how this information might be extracted from a page of text. Here, we dissected literacy-trained convolutional neural networks (CNNs) to formulate hypotheses about how letters and their positions are extracted from visual strings, then tested them using fMRI and MEG. In CNNs, literacy training led to the emergence of units with selectivity for words, particularly the trained script relative to untrained scripts. This finding is akin to the formation of a Visual Word Form Area (VWFA) (C. I. Baker et al., 2007; Cohen et al., 2002). The main advance here is that we clarify how the firing of these units collectively encodes a written word. We first show that their response profile fits an approximate letter x ordinal position code, with position being encoded as an approximate number of letters relative to both word edges. We then used a neuro-physiological approach to identify their preferred letter(s), their response to strings that orthogonally vary letter position, spacing, and word position, and their connections through earlier layers.

The outcome is a precise mechanistic hypothesis about the neural circuitry for invariant word recognition. This “space-bigram” model can be seen as a reconciliation of two previous proposals: letter bigrams and letter-position coding. In agreement with the letter-bigram model, IT neurons encode frequent character pairs – except that one of those characters is a space (which is indeed the most frequent character in text). As a result, neurons end up responding selectively to a single or a few letters at a given ordinal position from either word beginning or word ending— an approximate ordinal code, compatible with recent psychophysical and intracranial recordings (Agrawal et al., 2020; McCloskey et al., 2013; Woolnough et al., 2020). This code is extracted by a feedforward hierarchy of neurons with increasingly larger retinotopic receptive fields, jointly sensitive to the high-frequency shapes that make up letters and to the low-frequency patterns that signal word boundaries (see Schubert et al., 2021 for a similar finding). Thus, the ordinal positions of every letter in a word can all be extracted in a fast, parallel, feedforward manner, unlike previous models that relied on hypothetical mechanisms of serial processing or temporal coding (Norris, 2013; Whitney, 2001). Also, the shift from position-dependent to position-independent coding is progressive, and a few units in later layers of the network continue to respond to their preferred letters at specific retinotopic positions, thereby accounting for the ability to decode stimulus position in VWFA (Rauschecker et al., 2012).

A thorough test of the model will require high-density neuronal recordings in human VWFA and connected sites, which may not occur until one or two decades. Meanwhile, we tested the proposed transition to an ordinal letter code by using sophisticated, yet indirect methods of decoding and representational similarity analyses applied to human MEG and 7T fMRI signals. Consistent with a hierarchy of increasingly invariant stages, letter identity could best be decoded from the later layers of the model and from anterior regions of the ventral visual pathway (LOC and VWFA). Crucially, ordinal letter position was encoded invariantly for retinotopic location only in VWFA and at a rather late time window (220 ms post-stimulus). Both location and timing were in excellent agreement with prior evidence from fMRI (Dehaene et al., 2004), MEG (Pammer et al., 2004) and intracranial recordings (Thesen et al., 2012; Woolnough et al., 2020).

The proposed neural code can also explain several prior findings in the neuropsychology of reading. Psychophysical studies show that edge letters are crucial for efficient word recognition, while middle letters can be partially displaced or transposed (Xiong et al., 2019). This phenomenon, popularly known as the Cmabridge Effect (Grainger & Whitney, 2004), arises here because the letter position detectors exhibit sharp ordinal sensitivity for edge letters and increasingly broader tuning for middle letter positions (see figure 2C) (Gomez et al., 2008). This coding scheme can also explain the transposition errors of patients with letter-position developmental dyslexia (Friedmann & Gvion, 2001), the position-preserving perseveration errors of a patient with acquired alexia (McCloskey et al., 2013), and the existence of developmental attentional dyslexia, where readers experience the migration of letters from neighboring words while preserving their ordinal positions (Friedmann et al., 2010).

Overall, we delineate the stages of orthographic processing that lead to invariant visual word recognition. Beyond reading, the proposed hierarchical scheme for moving from retinotopic to relative-position codes readily extends to the recognition of the configuration of parts within an object, as studied in the macaque monkey (C. Baker et al., 2002). The same mechanism also allows to encode the position of parts, not only relative to left/right, but also to top/bottom, which is required for Chinese, mathematical or musical notations. Finally, while we focused entirely on the endpoint of learning, the present work could easily be extended to study the developmental emergence of letter position codes in both models and children (Dehaene-Lambertz et al., 2018).

## Methods

### Model architecture and training

Among the many available convolutional neural networks that can predict neural responses along the ventral visual pathway, we chose CORnet-Z architecture for two reasons: 1) It has a modular structure that resembles the stages of processing in the visual cortex (V1, V2, V4, IT, avgpool IT or H, output), 2) It has fewer parameters, thus lowering the training time while achieving high levels of accuracy on synthetic word datasets (Hannagan et al., 2021). Similar to our previous work, we first trained this network on the ImageNet dataset (phase 1), which contains ∼1.3 million images across 1000 categories. This was considered an illiterate network, which encodes the visual properties of objects but not text. Next, we extended the number of output nodes to 2000 (1000 images + 1000 words), with full connectivity to H layer units, and retrained the entire network jointly on ImageNet and a synthetic word dataset (phase 2), which also contained 1.3 million images of 1000 words. This was considered a literate network.

We trained separate networks in 5 different languages: French, English, Chinese, Telugu, and Malayalam. To investigate the mechanisms of bilingualism (Zhan et al., 2023), two additional networks were trained on bilingual stimuli: English + Chinese, and English + French. To estimate the variability across training sessions, each literate network was trained starting from the same five instances of illiterate networks. Thus, there were a total of 35 literate and 5 illiterate networks. Pytorch libraries were used to train these networks with stochastic gradient descent on a categorical cross-entropy loss. The learning rate (initial value = 0.01) was scheduled to decrease linearly with a step size of 10 and a default gamma value of 0.1. Phase 1 training lasted for 50 epochs and phase 2 training for another 30 epochs. The classification accuracy did not improve further with more epochs.

### Stimuli

To improve network performance, ImageNet images were transformed using standard operations such as “RandomResizedCrop” and “Normalize”. The images were of dimension 224x224x3. To avoid cropping out some letters in the word dataset, the default scale parameter of RandomResizedCrop was changed such that 90% of the original image was retained. For fair comparisons, other operations such as flipping were not performed on the Imagenet dataset as the same operation would create mirror words in the Word dataset, which is not typical in reading.

The English and French words included frequent words of length between 3-8 letters. The Chinese words were 1-2 characters long. The Telugu and Malayalam words were 1-4 characters long, which approximates the physical length of the chosen English/French words. The synthetic dataset comprised 1300 stimuli per word for training and 50 stimuli per word for testing. These variants were created by varying position (-50 to +50 along the horizontal axis, and -30 to +30 along the vertical axis), size (30 to 70 pts), fonts, and case (for English and French). For each language, 5 different fonts were chosen: 2 for the train set, i.e. Arial and Times New Roman for English and French; FangSong and Yahei for Chinese; Nirmala and NotoSansTelugu for Telugu; and Arima and AnekMalayalam for Malayalam; and another 3 fonts for the test set, i.e. Comic Sans, Courier, and Calibri for English and French; Kaiti, simhei, and simsum for Chinese; NotoSerifTelugu, TenaliRamakrishna, and TiroTelugu for Telugu; and Nirmala, NotoSansMalayalam, and NotoSerifMalayalam for Malayalam.

The bigrams stimuli used in the dissimilarity analysis (Figure 1C-D) were taken from earlier studies (Agrawal et al., 2019, 2020).

### Identification of word selective units

Similar to fMRI localizer analysis, a unit was identified as word selective if its responses to words were greater than the responses to nonword categories: faces, houses, bodies, and tools by 3 standard deviations. The body and house images were taken from the ImageNet dataset. For tools, we used the “ALET” tools dataset, and Face images were taken from the “Caltech Faces 1999” dataset. We randomly chose 400-word stimuli and 100 images each from the other categories for identifying category-selective units.

### Dissimilarity measure

For each layer, we vectorized the activation values, and estimated the pair-wise dissimilarity value using correlation metric i.e., d = 1-r. where r is the correlation coefficient between any two activation vectors.

### Letter selectivity

For each word-selective unit, its tuning profile across letters was estimated by identifying the letter that evoked maximum response at either of the 8 possible positions. First, we obtained 26x8 = 208 responses, followed by the max operation across positions, and were further sorted to identify the most and least preferred letter for a given unit.

### Activation maximization

We used the Lucent toolbox (https://github.com/greentfrapp/lucent) to generate images that maximally activate a given unit. Given the convolutional structure of the early stages of the network, all units within a given channel have the same features but at different spatial locations. Thus, we only estimated the preferred input for the unit whose receptive field was at the center of the image. We generated images that maximized activation in both positive and negative directions with a threshold of 1000 iterations. For ease of visualization, only the images generated along the positive directions are shown.

### fMRI data

#### Participants

A total of 16 adult subjects (aged b/w 18-40 years) who were native French speakers participated in this study. They were right-handed and had normal or corrected-to-normal vision with no history of any psychiatric disorders or reading difficulties. Participants provided written informed consent for the fMRI study and received monetary compensation. The study was approved by the local ethics committee in the NeuroSpin Centre (CPP 100055) and was conducted following the Declaration of Helsinki.

#### Stimuli

The functional localizer block consisted of various stimuli, including French words, objects, scrambled words, and scrambled objects. For each run, 12 unique images were randomly chosen from a pool of available images. The scrambled images were created by scrambling the phase of the Fourier-transformed images and then reconstructing them using the inverse Fourier transform.

The event block comprised 36 unique stimuli, which consisted of three-letter strings (e.g., OXX, XOX) with specific characteristics. Each string contained one frequent letter (e.g. O) and two repeated rare letters (e.g. X). We specifically selected three letter pairs for this purpose: OX, TB, and EQ, with the first letter in each pair being frequent and the second being rare in French, the language of our participants. These letter pairs were also chosen based on their visual dissimilarity and low bigram frequency. The design was founded upon the hypothesis that the frequent letter would cause activation in putative letter-selective units, while variations in its absolute location on the screen and in its relative location within the string would allow us to establish the relative or positional nature of its neural code. A 3x4 design of the letter strings was used to dissociate the absolute and ordinal positions of the frequent letters (Figure S1). Absolute word position was varied by shifting the strings by one letter at a time, ranging from completely left to completely right of the fixation dot (four levels of word position, resulting in six levels of absolute letter position). At each word position, the odd letter could appear in each of three positions within the 3-letter string (left, center, or right).

The stimuli were presented on a BOLD screen (Cambridge Research Systems, Rochester, UK), a 32-inch MRI-compatible LCD screen with a resolution of 1920 × 1080 pixels. The screen had a refresh rate of 120 Hz and was positioned at the head-end of the scanner bore. Participants viewed the stimuli through a mirror attached to the head coil.

#### Task design

The localizer run comprised 14-s mini blocks in which a total of 14 images were presented for 0.8 s with an inter-stimulus interval of 0.2 s. There were 12 unique stimuli and 2 of them repeated at random time points, corresponding to targets for the one-back task. Following each block of stimuli, a blank screen with a fixation cross was presented for 6 seconds. Each block (words, objects, and their corresponding scrambled versions) was repeated three times within each run. There were two runs of the localizer.

In the event-related runs, each of the 36 stimuli was displayed for 0.2 seconds, followed by a blank screen lasting 2.8 seconds. Within each run, each stimulus was repeated five times. To actively engage the participants, twenty task trials were also included, during which subjects were required to respond with a button press when all the letters in a stimulus were identical (e.g., XXX, BBB, QQQ). Additionally, to introduce variability in the inter-stimulus interval, 10% of the trials (n = 20) did not present any stimulus. This helped to jitter the timing between stimuli. There was a total of 4 runs, 660 s long, and each run started and ended with a blank screen displaying a fixation cross for 4s. The high-resolution MP2RAGE anatomical images were obtained in the middle of the scan session i.e., after two event-related and one localizer run. The subjects were instructed to close their eyes and relax during the anatomical scan.

#### Data acquisition

The Brain images were acquired using a 7-T Magnetom scanner (Siemens, Erlangen, Germany) with an a1Tx/32Rx head coil (Nova Medical, Wilmington, USA) at the NeuroSpin Centre of the French Alternative Energies and Atomic Energy Commission. Dielectric pads were placed around the ear to reduce the signal dropout around the anterior Ventral Occipital Temporal cortex.

Functional data were acquired with a two-dimensional (2D) gradient-echo echo-planar imaging (EPI) sequence using the following parameters: repetition time (TR) = 2000 ms, echo time (TE) = 21ms, voxel size = 1.2 mm isotropic, multiband acceleration factor = 2; encoding direction: anterior to posterior, iPAT = 3, flip angle = 75, partial Fourier = 6/8, bandwidth = 1488 Hz per pixel, echo spacing = 0.78 ms, number of slices = 70, no gap, reference scan mode: GRE, MB Leak Block kernel: off, fat suppression enabled. To correct for EPI distortion, a five-volume functional run with the same parameters except for the opposite phase encoding direction (posterior to anterior) was acquired immediately before each task run. Participants were instructed not to move between these two runs. Manual interactive shimming of the B0 field was performed for all participants. The system voltage was set at 250 V for all sessions, and the fat suppression was decreased to ensure that the specific absorption rate did not surpass 62% for all functional runs. High-resolution MP2RAGE anatomical images were obtained in the middle of the session with the following parameters: resolution = 0.65 mm isotropic, TR = 5000 ms, TE = 2.51 ms, TI1/TI2 = 900/2750 ms, flip angles = 5/3, iPAT = 2, bandwidth = 250 Hz/Px, echo spacing = 7ms.

#### Data preprocessing

The raw functional data underwent distortion correction using FSL topup (https://fsl.fmrib.ox.ac.uk/fsl/fslwiki/topup). This distortion-corrected data was further processed using the SPM12 toolbox (https://www.fil.ion.ucl.ac.uk/spm/software/spm12). The functional images were realigned, slice-time corrected, co-registered with anatomical images, segmented, and finally normalized to the MNI305 anatomical template. All SPM parameters were set to default and the voxel size after normalization was set to 1.2x1.2x1.2 mm^3^. Before normalization, the data were denoised using GLMdenoise (Kay et al., 2013), which is known to improve signal-to-noise ratio.

The preprocessed data were concatenated across all runs and modeled using a generalized linear model (GLM) in SPM using the default parameters. For the event-related run, the t-values were estimated for each condition and were used for further analysis instead of beta values.

#### ROI definitions

The regions of interest along the ventral visual pathway were identified by contrasting the activation between conditions from the localizer run. Early visual areas were identified by contrasting scrambled objects with fixation cross. This region was further parsed into V1-V3, and V4 using the anatomical mask from the SPM anatomy toolbox (Eickhoff et al., 2005). Higher visual area (LOC) was identified as a region that responded more to objects than scrambled objects. The voxels in the LOC region were restricted to the Inferior Temporal Gyrus, Inferior Occipital Gyrus, and Middle Occipital Gyrus. These anatomical regions were obtained from Tissue Probability Map (TPM) labels in SPM 12. VWFA was identified as a region in the left occipital temporal sulcus that responded more to words than scrambled words. For each contrast, a voxel-level threshold of p < 0.001 (uncorrected) was used to obtain a contiguous region.

### MEG data

#### Participants

A total of 18 adult subjects (aged b/w 18-40 years) who were native French speakers participated in this study. They were right-handed and had normal vision with no history of any psychiatric disorders or reading difficulties. Participants provided written informed consent for the MEG study and received monetary compensation. One subject was excluded from this study due to excessive head movements.

#### Stimuli

The stimuli were identical to the fMRI study

#### Task design

As in the fMRI, a series of letter string stimuli were presented to the participants, who had to respond with a button press if all the letters were identical. Each stimulus was shown for 200ms followed by a 300ms blank screen with a fixation cross. In a run, each stimulus was repeated 15 times, plus 60 target trials. The study comprised three such runs in total.

#### Data acquisition

Participants performed the tasks while sitting inside an electromagnetically shielded room. The magnetic component of their brain activity was recorded with a 306-channel, whole-head MEG by Elekta Neuromag (Helsinki, Finland). 102 triplets, each comprising one magnetometer and two orthogonal planar gradiometers composed MEG helmet. The brain signals were acquired at a sampling rate of 1000 Hz with a hardware highpass filter at 0.03 Hz. Eye movements and heartbeats were monitored with vertical and horizontal electrooculograms (EOGs) and electrocardiograms (ECGs). Head position inside the helmet was measured at the beginning of each run with an isotrack Polhemus Inc. system from the location of four coils placed over frontal and mastoidian skull areas.

#### Data preprocessing

The MEG data underwent several preprocessing steps. Initially, the data was inspected to identify any sensors showing excessive noise or poor signal quality. This was performed using a custom script in which sensors displaying deviations from the median variance across time by more than 6 standard deviations were identified as deviant sensors and excluded from further analysis.

To remove environmental noise and artifacts related to head movements, Maxwell filtering was applied using the MaxFilter tool in MNE-Python. This step compensated for head position changes during the recording session by using the continuous head position indicator (cHPI) coil signals. The data were processed with signal space separation (SSS) to suppress external interference and enhance the signal-to-noise ratio. A finite impulse response bandpass filter was then applied to the data to focus on the frequency range of interest (1-40 Hz) and eliminate unwanted noise.

Signal space projection (SSP) techniques were employed to further improve data quality and reduce artifacts. SSP vectors were computed by identifying signal components, such as eye blinks or heartbeats. These SSP vectors were then projected out from the MEG data to remove the corresponding artifacts.

#### Eye tracking

During the MEG acquisition, participants were instructed to fixate on the central cross while the items were flashed. Their gaze was monitored online using EyeLink 1000 eye-tracker device (SR research). Eye-tracking data was collected for 16 out of 18 participants

#### Decoding analysis

The stimuli used in fMRI and MEG experiments varied along the following factors: absolute word position, absolute letter position, ordinal letter position, and letter identity. To analyze the neural signals corresponding to these factors, a linear classifier was used. Specifically, logistic regression with a “liblinear” solver was performed after scaling the data using “robustscalar” from the sci-kit learn package in Python. To avoid overfitting, a 3-fold cross-validation was performed using the MNE function “cross_val_multiscore”. For decoding the position information, the classifier was trained on the data from any two letter pairs, and tested on the third one. Similarly, for decoding letter identity, the classifier was trained on any two ordinal positions and tested on the remaining one. In fMRI, the voxels within each region were used as features, and in MEG, the channels were used as features with data averaged within 10ms time-bins.

#### Estimating neural dissimilarity

In fMRI, for a given ROI and subject, pair-wise dissimilarity between a stimulus pair was computed using correlation distance metric (i.e., 1 – r); where r is the correlation coefficient between the activity pattern across voxels. Similarily, in MEG, the signal across each channel was first baseline corrected using MNE function “mne.baseline.rescale” by subtracting the mean and dividing by the standard deviation of the baseline values (zscore). The correlation distance was then estimated using channels as features

## Notes

### Competing Interest Statement

The authors have declared no competing interest.

