## Supplementary material for "Dissecting the neuronal mechanisms of invariant word recognition": Supplemetary text

### Supplementary materials

- Supplementary Figure 1: Stimuli used in Brain imaging studies
- Supplementary Figure 2: Behavioral Task Performance
- Supplementary Figure 3: Representational similarity space predicted by literate CNN
- Supplementary Figure 4: Examples of units with mixed selectivity
- Supplementary Section 1: Emergence of ordinal position coding units in literate CNN
- Supplementary Section 2: Comparative analysis of CNN, fMRI, and MEG
- Supplementary Section 3: Searchlight analysis

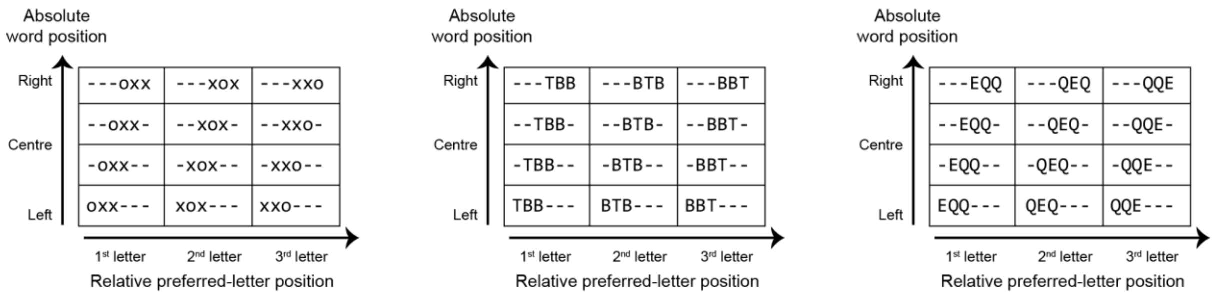


**Figure S1: Stimuli used in brain imaging studies.** Schematic of the stimuli used to dissociate position coding. The word position varies across rows, and the preferred letter position within a string varies across columns. '-' represents blank space. Overall, there are 36 unique stimuli.

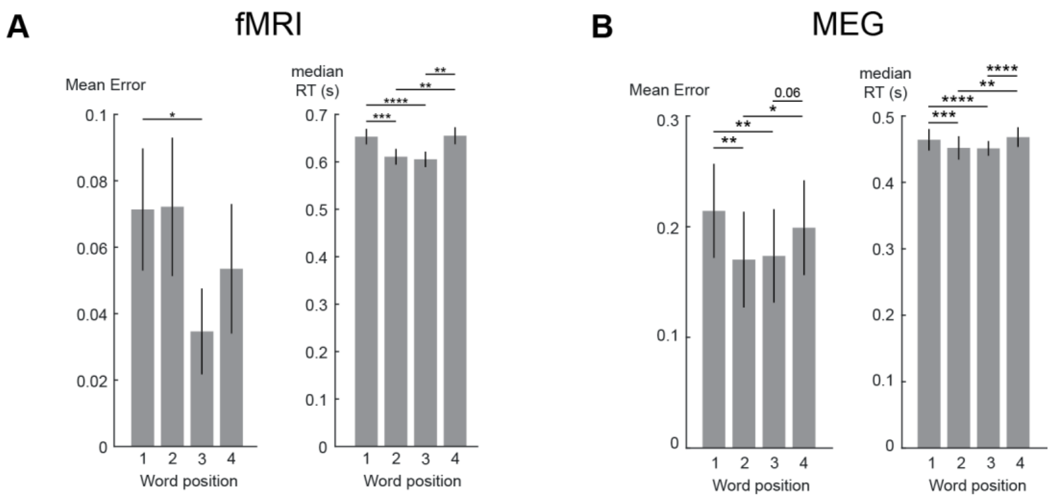


**Figure S2: Behavioral task Performance.**

(A) During fMRI data acquisition, subjects were instructed to respond with a button press when all the letters in a stimulus were identical. The stimuli could appear at any word position during task trials. Subjects exhibited low error (left) and faster response times (right) when the stimuli were presented at the fovea. Each run included 10% task trials. Error bars indicate standard deviation across subjects and the asterisks indicate statistical significance using paired t-test (* $P < 0.05$, ** $p < 0.005$, *** $p < 0.0005$, etc).

(B) Same as (A) but for MEG data. Faster stimulus presentation with an inter-trial interval of 300 ms resulted in a speed-accuracy trade-off: in MEG, compared to fMRI, subjects were less accurate but responded faster.

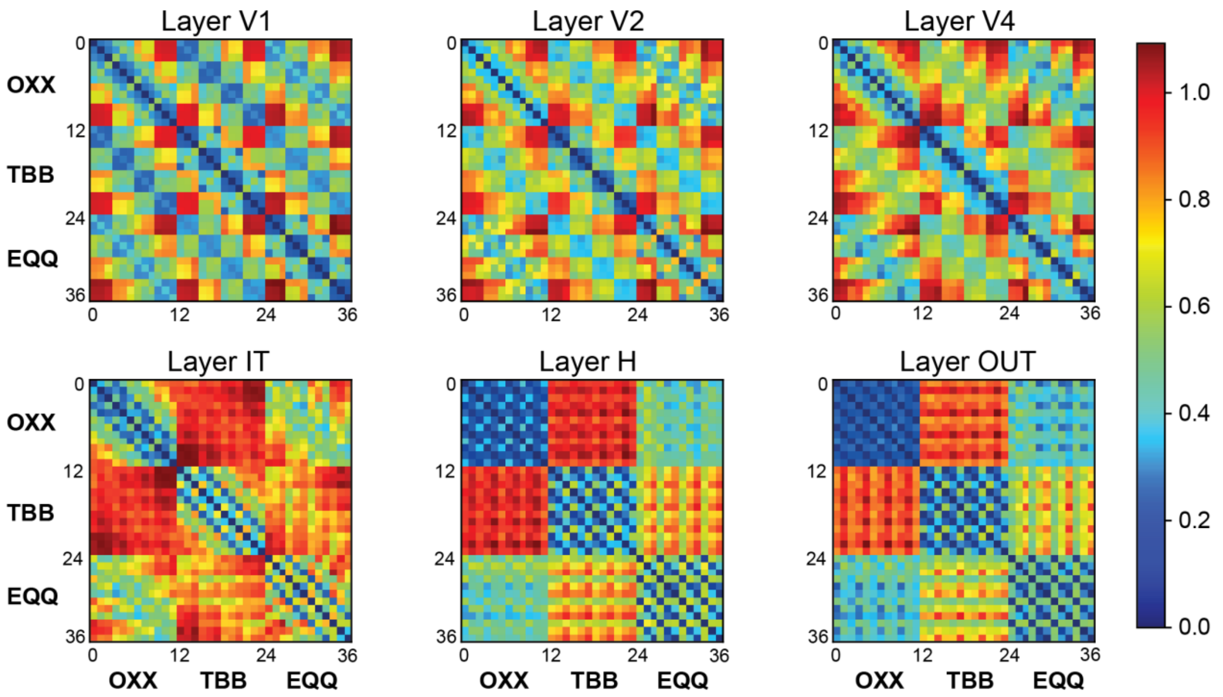


**Figure S3: Representational similarity space predicted by literate CNN.** Representation Dissimilarity Matrices (RDMs) predicted by different layers of the literate network across the 36 stimuli that were used in the brain-imaging experiments. The correlation distance metric (1-r) is plotted according to the color scale at right. In the early layers, word position and absolute letter positions are encoded irrespective of the specific letter identities. The IT layer demonstrates partial invariance to word position and encodes letter identity. Complete position invariance is observed from the H layer.

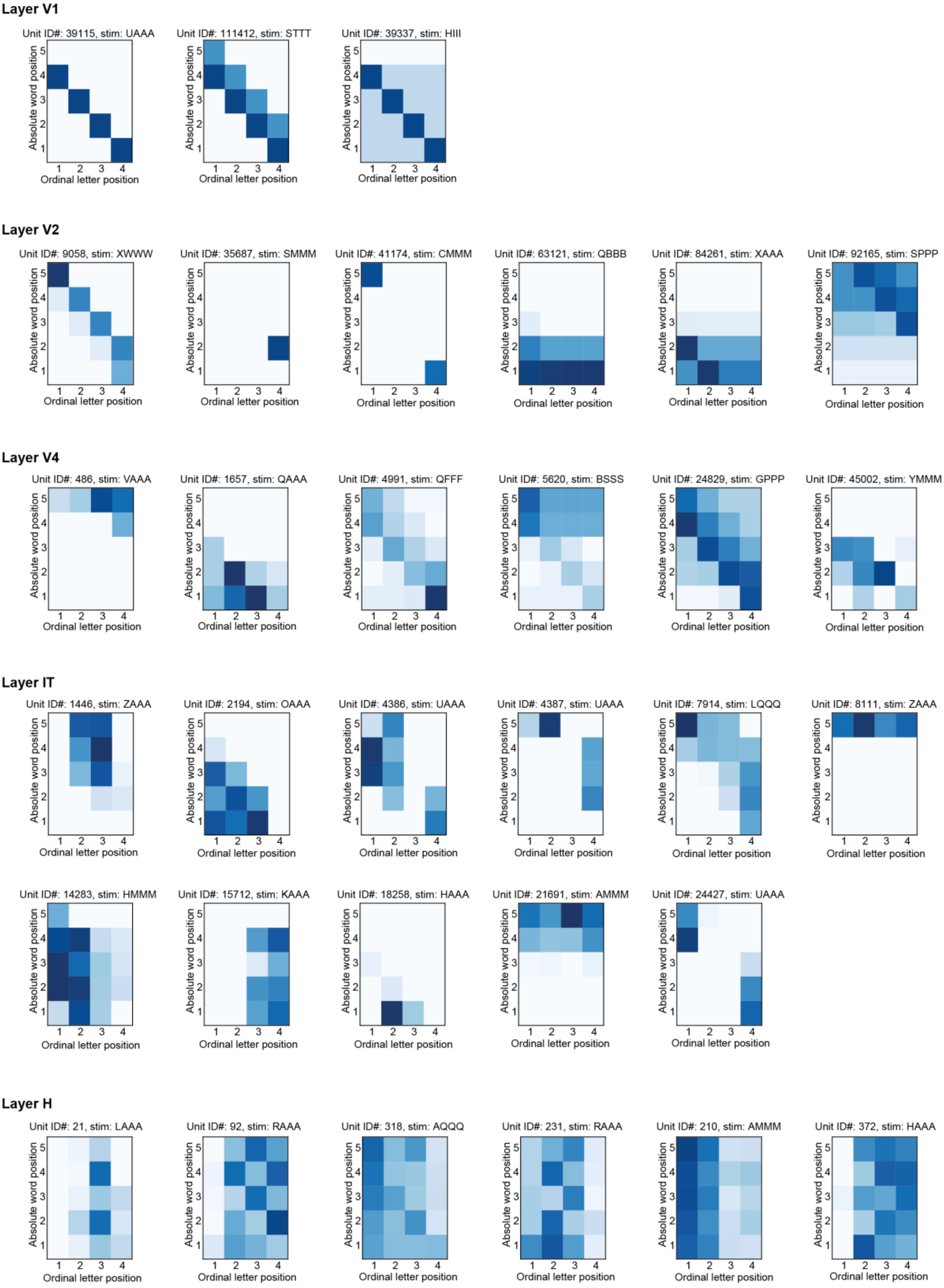


**Figure S4: Examples of units with mixed selectivity.** Diversity of response profile across word-selective units from different layers of the literate network.

**Supplementary section 1: Emergence of ordinal position coding units in literate CNN**

The response profile of units across different layers of the literate network suggested that ordinal position coding in higher layers (e.g., IT) arises because those units pool across blank-space coding units in the earlier layer (e.g., V4). To test this hypothesis, we investigated in detail the input connection profile of IT units. Our goal was to characterize, mechanistically, how each IT unit got its selectivity by visualizing its strongest V4 input units and understanding, in turn, how the cells were tuned. We report here the example of two IT units (figure S5).

Each layer in a CNN undergoes two stages of processing: Convolution and Max Pooling. Here, the convolution filters have a kernel size of 3x3, along with padding and a stride of 1. Consequently, the dimensions of the output matrix after the convolution operation remain unchanged. The Maxpool2d operation also employs a 3x3 kernel with a stride of 2, which reduces the dimension of the output matrix to half of the input. In our network, the dimensions of the V4 layer are 14x14x256. In this context, 14x14 represents the spatial position, and there are 256 distinct features extracted at any given spatial position (256 "filters"). Each IT unit is therefore connected to the V4 layer through 256 weight matrices, each of which has a dimension of 3x3. Since these weights are applied to outputs that have undergone two stages of processing, the effective receptive field of an IT unit relative to the V4 layer is 5x5.

To characterize the functional connectivity of a given IT unit from V4 units, we began by summing the weights of each convolution filter, resulting in a 256-length vector. We opted for this summation approach instead of using L2norm or any other absolute measure, primarily because the ReLU operation nullifies all negative responses, ensuring that negative weights only dampen activity. Next, within the 5x5 receptive field of an IT unit, we identified word-selective units and assessed their responses to the preferred stimulus of the chosen IT unit. The stimulus set consisted of 20 stimuli created by combining a single preferred letter and 3 copies of the least preferred letter at various ordinal positions (refer to figure 3A and the main text for details). For each word-selective unit in V4, the maximum response across those 20 images was recorded. To obtain a single input activation value for each of the 256 V4 filters, we performed the max operation on all word-selective units for a given filter, resulting in another set of 256-length vectors.

The product of the input V4 activation with the weight vector enabled us to distinguish between features that either activate or inhibit the response of the selected IT unit. For each of two IT units, Figure S5 shows the top two V4-to-IT connectivity filters that most significantly contribute to its response. Our observations indicate that each IT cells receives two types of V4 inputs: (1) V4 cells that are already highly selective to one or a few letters, but with a broad retinotopic receptive field; and (2) V4 cells that typically have broader selectivity for several letter identities, but care about blank spaces too and therefore begin to exhibit ordinal coding (see figure S5).

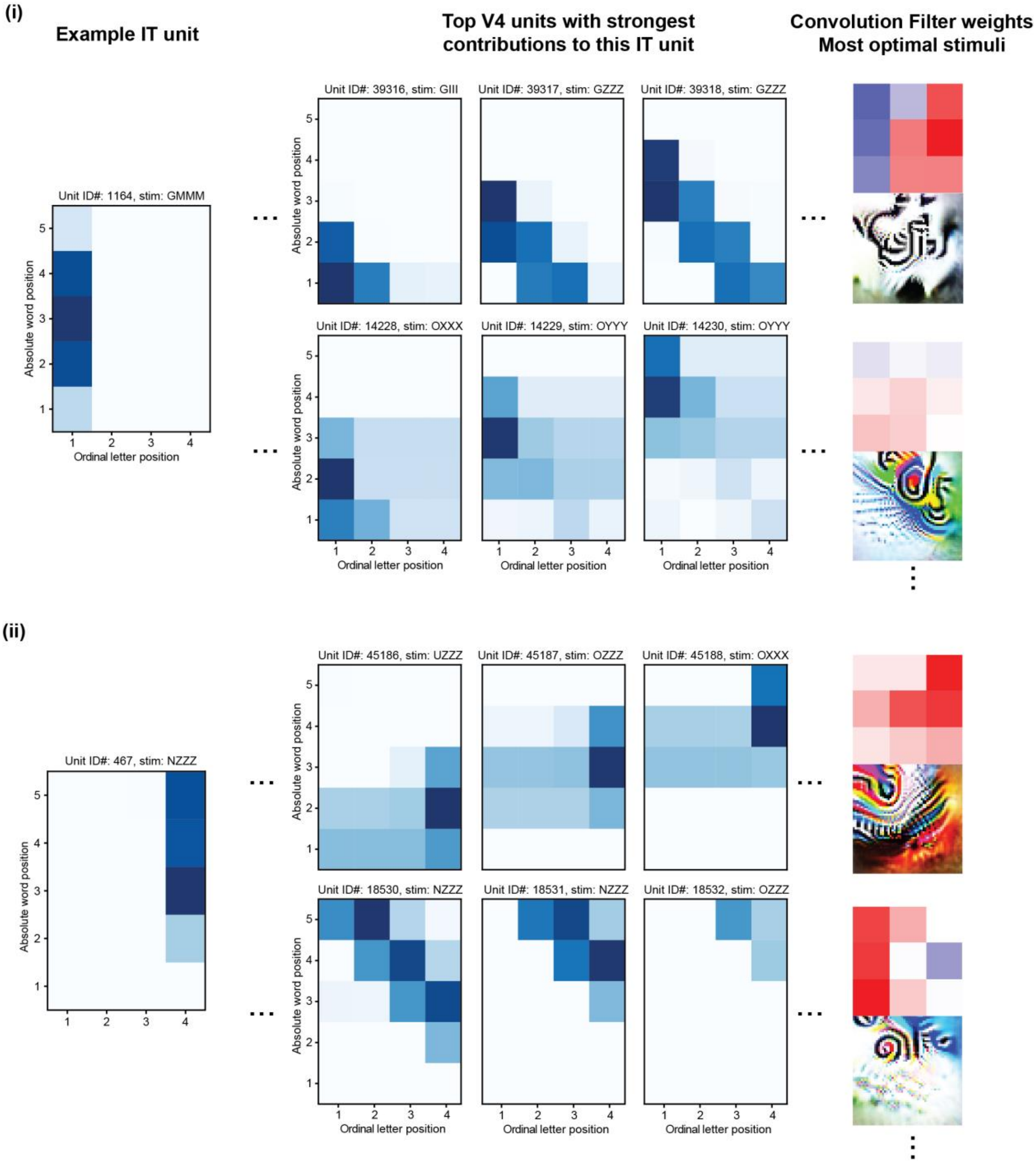


**Figure S5: Functional connectivity between V4 and IT units.** (left) response profile of an example IT units that encodes ordinal position; (middle) Response profile of word selective units in the V4 layer to their own preferred stimuli that lie within the receptive field of the specific IT unit; (right) Top-2 convolution filter weights that connect V4 and IT layer and the stimuli that would maximally activate a given filter generated using activation-maximization method.

To further probe this, we examined the responses of these units to single letters presented independently at each of the 8 spatial positions (Figure S6). Our observations confirmed that units with retinotopic coding exhibit preferential responses to a select few letters within their receptive field, while blank space coding units respond to a broader range of letters (Figure S6).

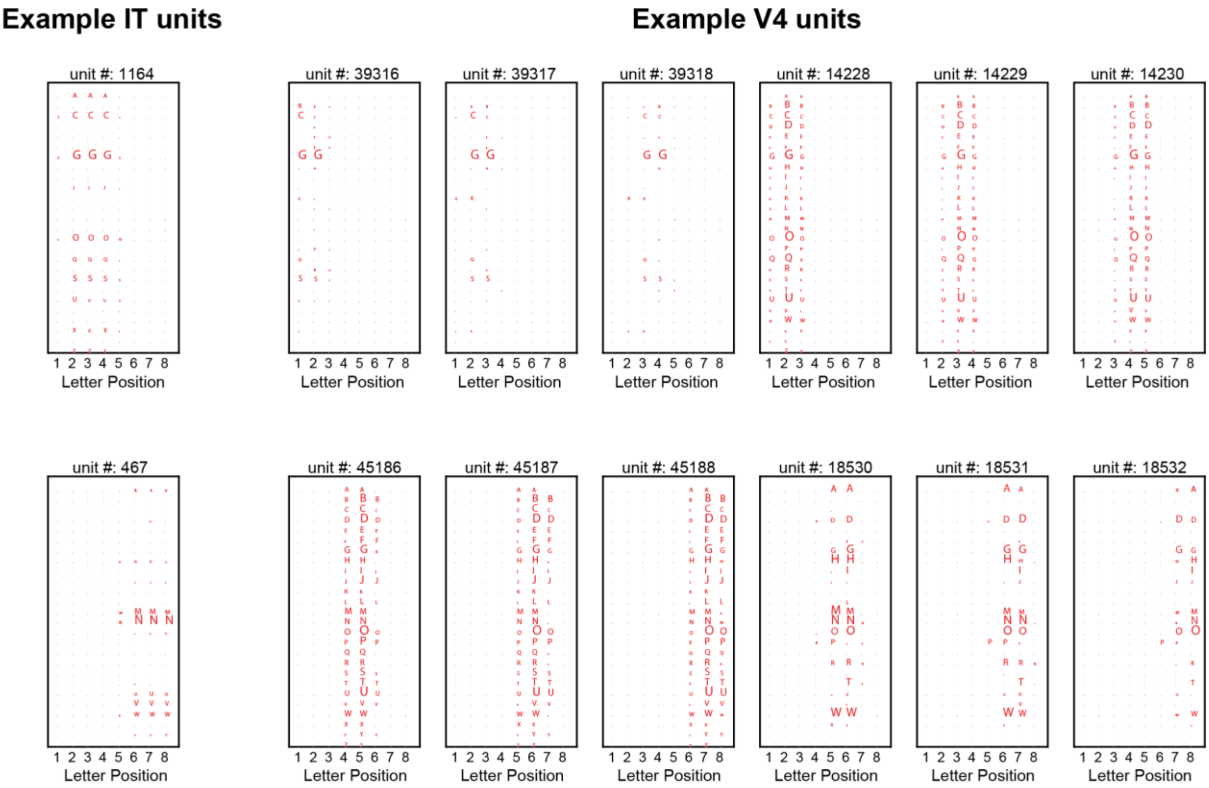


**Figure S6: Response of V4 and IT units to individual letters.** For the same 2 IT units and 6 V4 units as in figure S5, the figure shows the response to single letters presented independently at each of 8 different spatial positions. The size of the letter indicates the magnitude of normalized responses.

Together, figures S5 and S6 show that higher-level units acquire their broad receptive fields and, simultaneously, their sensitivity to specific letters at a given ordinal position, through the pooling of units at the immediately preceding layer that are (1) less spatially invariant, but (2) already sensitive to some letters and the presence of a space at some distance to the left or to right. The pooling mechanism simultaneously abstract away from retinotopic information while refining each unit's letter specificity and ordinal coding.

### Emergence of blank space coding

Having established the link between space coding and ordinal position units in the higher layers of the literate CNN, we next investigated the type of inputs from the V1 layer that leads to the formation of space coding units in the V2 layer. Since the V1 layer has very few word-selective units, we could not perform the same analysis as just described. Instead, we initially visualized the weights of the V1 layer to gain insights into which features in the image would elicit the maximum response (Figure S7). To ensure all filter weights fell within the range [0, 255], any negative values were adjusted by adding the minimum value and normalizing the maximum value. This analysis revealed a classical array of early visual features, including Gabor-like filters and color boundaries.

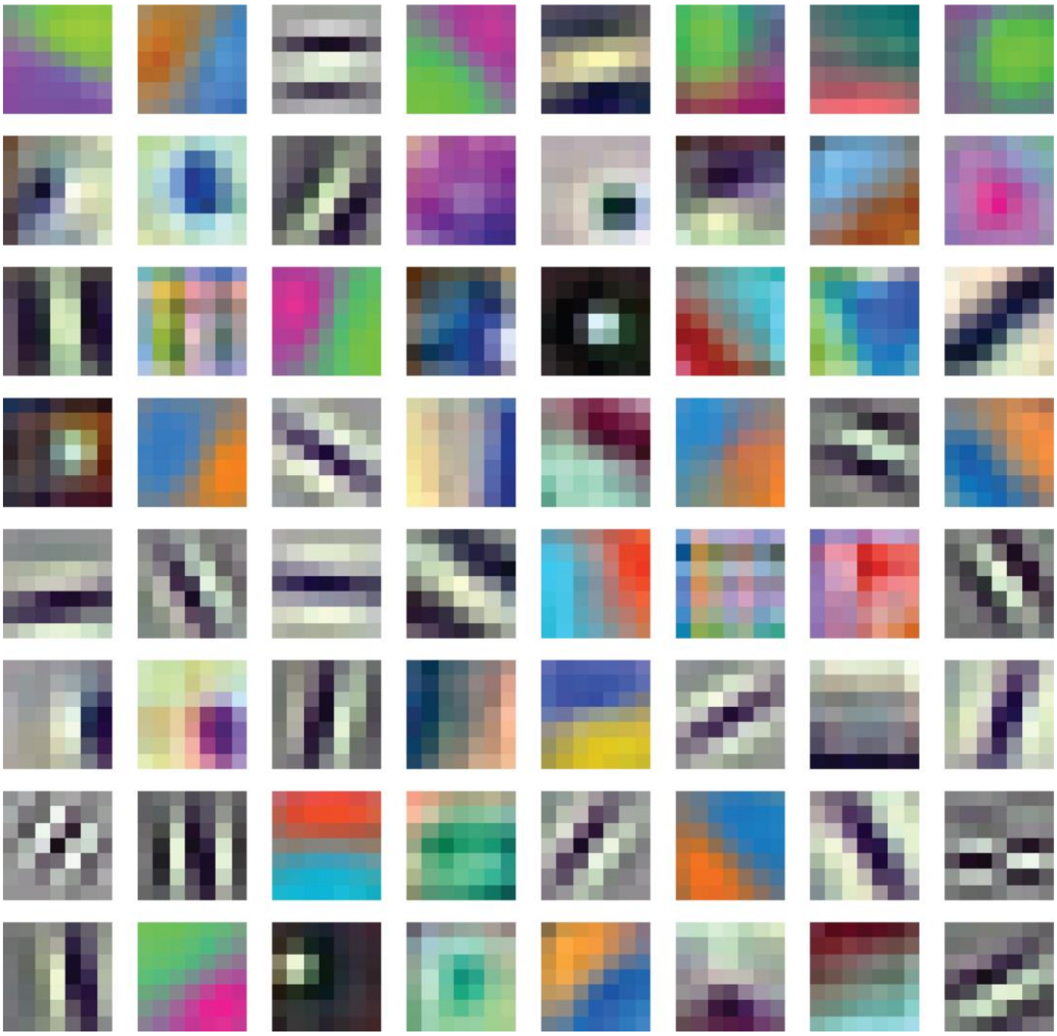

**Figure S7: Visualization of V1 filters.** The V1 units comprise 64 distinct filters, each of size 7x7, whose weights are modified during training. Each filter represents the type of input image that would evoke a maximum response.

Following this, for any given V2 unit, as in our earlier approach, we summed the weights of its convolution filter from V1, and estimated the input activation vector across all units within its receptive field. For each V2 unit, we identified the top-5 inputs from V1 that resulted from the product of summed convolutional filter weights and V1 unit activations (see figure S8 for 4 example units). Interestingly, we observed that the retinotopic units in V2 (bottom two example units in figure S8) are primarily driven by V1 units equipped with ridge detector filters. Such V1 units are ideally suited to detect individual character parts, since they respond to high-spatial-frequency features such as oriented bars or curves, whenever there is an intensity change from white to black to white again in a close sequence. Such features are characteristic of the fine strokes and curves that make up characters in text. By contrast, we found that space coding units (top two example units in figure S8) are predominantly influenced by V1 units with edge-detection filters, i.e. low-spatial-frequency filters that encode the broader transitions between white and black pixels. Such filters are most suited to detect the overall shape or contour of a word and especially the location of its beginning and ending.

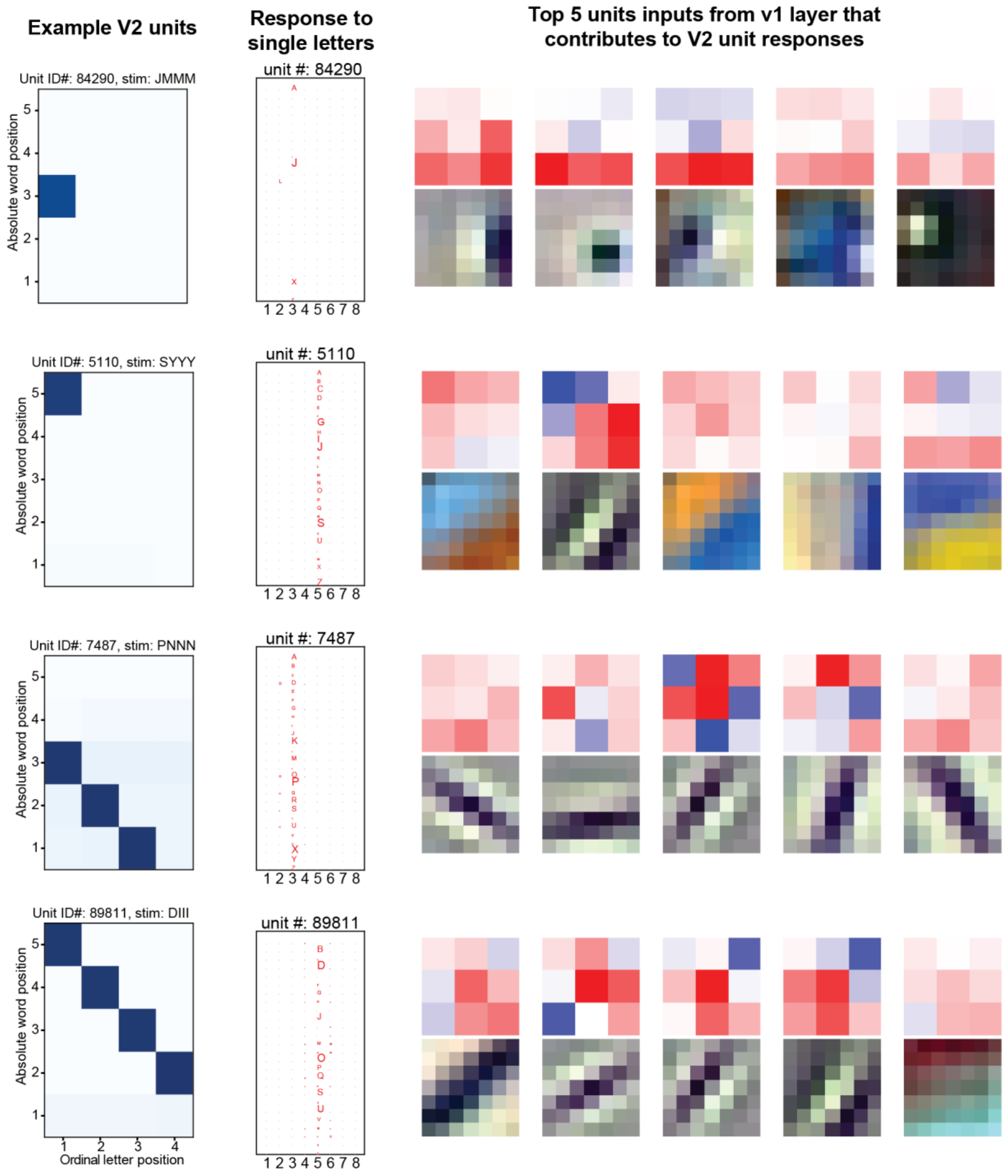

**Figure S8: Functional connectivity between V1 and V2 layers.** The figure shows the responses of four V2 units: two space coding units (top rows) and two retinotopic coding units (bottom rows). The units are characterized by their responses to 4-letter stimuli dissociating word position and ordinal letter position (left column; see figure 3A) and to individual letters at each of 8 retinotopic positions (second column). The right-hand side shows the top-5 convolutional filter weights between V1-V2 and corresponding features in the image that would elicit maximum response.

### Supplementary section 2: Comparative analysis of CNN, fMRI, and MEG

Representational similarity analysis provides a measurement tool by which the representational spaces can be compared across different branches of neuroscience such as fMRI, MEG, neurophysiological recordings or computational models [1,2]. Here, the comparison of representational dissimilarity matrices (RDMs) across different layers of the CNN, with different ROIs in fMRI, and different timesteps in MEG allowed us to investigate the existence of shared or distinct representations in different modalities and processing stages. These analyses provide insights into the information encoding and processing mechanisms underlying the observed brain activity.

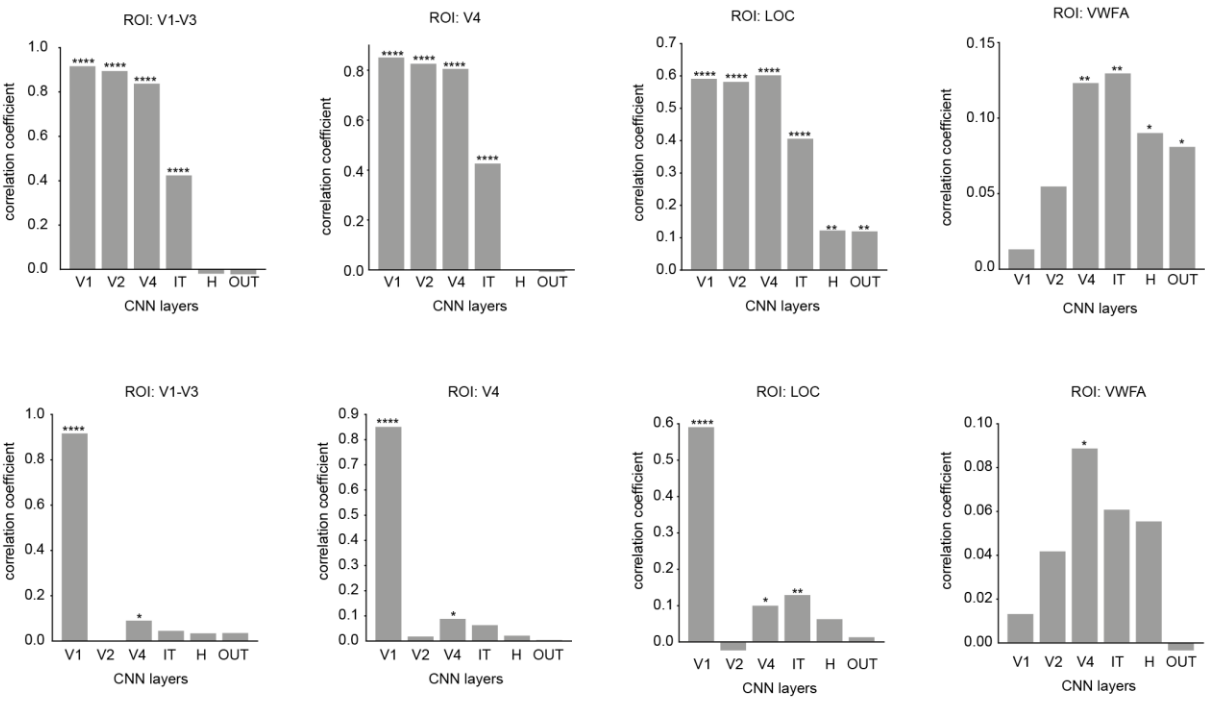


**Figure S9: Comparison of the representational spaces between fMRI and CNN**

(A) Each panel illustrates the correlation between the RDMs obtained from different layers of the network, represented on the X-axis, and those of a specific region of interest (ROI). The ROIs include V1-V3 (primary visual cortex), V4 (extrastriate visual cortex), LOC (lateral occipital complex), and VWFA (visual word form area). The high correlation values indicate the similarity of the representation space between the network layers and the activity within each ROI.

(B) Same correlation analysis as in (A), but after regressing the variability already contributed by the previous layers. This approach allows for a more focused examination of the unique contribution of each layer to the activity within the ROIs. By removing the shared variability, the correlation values provide insights into the specific relationships between the network layers and the targeted ROIs, independent of the influences from previous layers. Asterisks indicate statistical significance (* $P < 0.05$, ** $p < 0.005$, *** $p < 0.0005$, etc).

**Comparing CNN and MEG RDMs**

Following preprocessing of the MEG signal, we performed further analysis to explore the relationship between MEG activity and the layers of the CNN. First, the MEG activity was averaged across 10ms time windows and repetitions to obtain a representative measure of the response. Next, we computed Representation Dissimilarity Matrices (RDMs) using the correlation distance metric (1-r), considering all 306 channels of the MEG data.

We then correlated the MEG RDMs at each time window with the RDMs obtained from each layer of the CNN. Notably, the representation in the early layers of the CNN demonstrated significant correlations across all time points of the MEG signal, starting from 60ms. This finding provides validation that the low-level retinotopic properties are preserved throughout the word recognition process. In contrast, the representation in the later layers of the CNN exhibited significant correlations specifically with the later time points of the MEG signal (Figure S10A). This finding aligns with the results obtained from the decoding analysis (as depicted in Figure 6), further supporting the consistency of our findings. However, the reason behind the observed high correlation in the time window around 40ms remains unclear.

To examine the relationship between CNN layers and MEG RDMs while accounting for the influence of earlier layers, we conducted a correlation analysis similar to what is done in fMRI studies. By regressing the contribution of earlier layers, we were able to focus specifically on the unique contribution of each layer to the MEG RDMs. Remarkably, our analysis revealed a strong correlation between the activity in the V1 layer and the IT layer of the CNN with the MEG RDMs (Figure S10B). This indicates that these layers play a prominent role in shaping the observed patterns of neural representations captured by the MEG data.

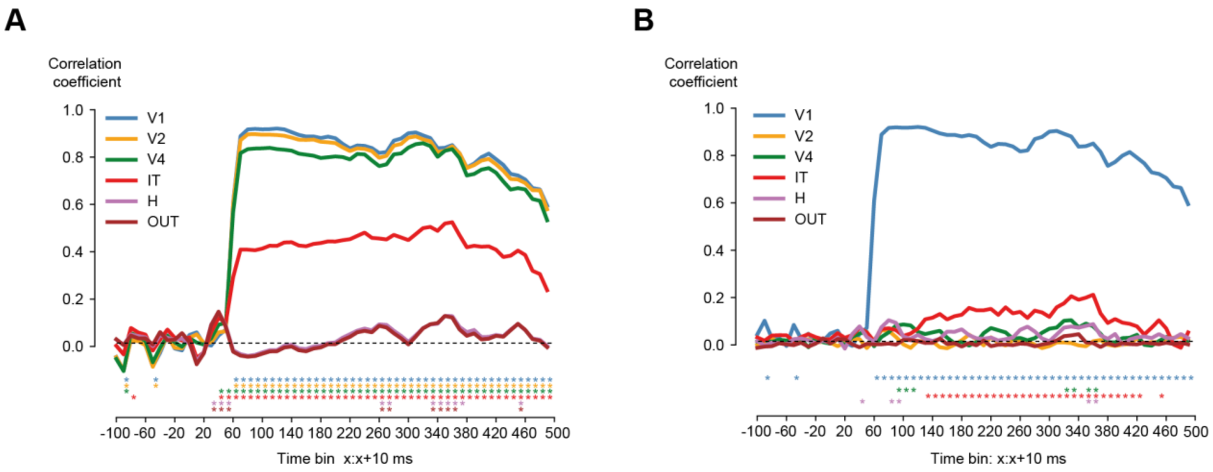


**Figure S10: Relationship between MEG and CNN**

(A) Correlation between MEG representation dissimilarity matrix (RDM) at each time point with different layers of the network. Asterisks indicate significant correlation ($p < 0.05$, spearman rank correlation).
(B) Same as (A) but after regressing out the contribution of earlier layers of the network.

**Comparing fMRI and MEG RDMs**

In line with the aforementioned comparison, we conducted a study examining the representation spaces of various regions in fMRI and MEG at different time intervals. Comparable to the initial layers of the cognitive network, the representation derived from early visual areas (V1-V3) displayed significant correlation across all time points of the MEG signal, commencing at 60ms. Additionally, each region made distinct contributions to the MEG RDMs.

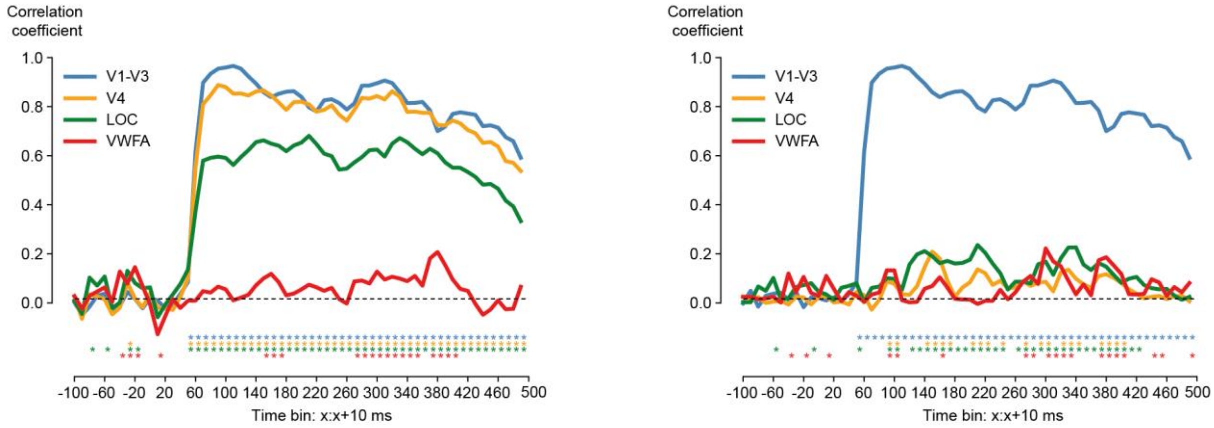


**Figure S11: Relationship between MEG and fMRI ROIs**

(A) Correlation between MEG representation dissimilarity matrix (RDM) at each time point with different regions of interest from fMRI data. Asterisks indicate significant correlation ($p < 0.05$, spearman rank correlation).

(B) Same as (A) but after regressing out the contribution of earlier regions.

### Supplementary section 3: Searchlight Analysis

To perform a more comprehensive and unbiased exploration of the neural correlates of the CNN layer in the whole brain, we performed a searchlight analysis. This involved examining each voxel within the brain and considering its surrounding neighbourhood of 343 voxels (7x7x7) to construct a representational dissimilarity matrix (RDM) using the correlation distance metric. We then correlated this RDM with different layers of the literate networks (Figure S12).

Additionally, to determine the unique contribution of each layer, we performed a step-wise regression analysis. This involved correlating the fMRI RDM with the CNN RDM, which was constructed after regressing the predictability of previous layers (Figure S13).

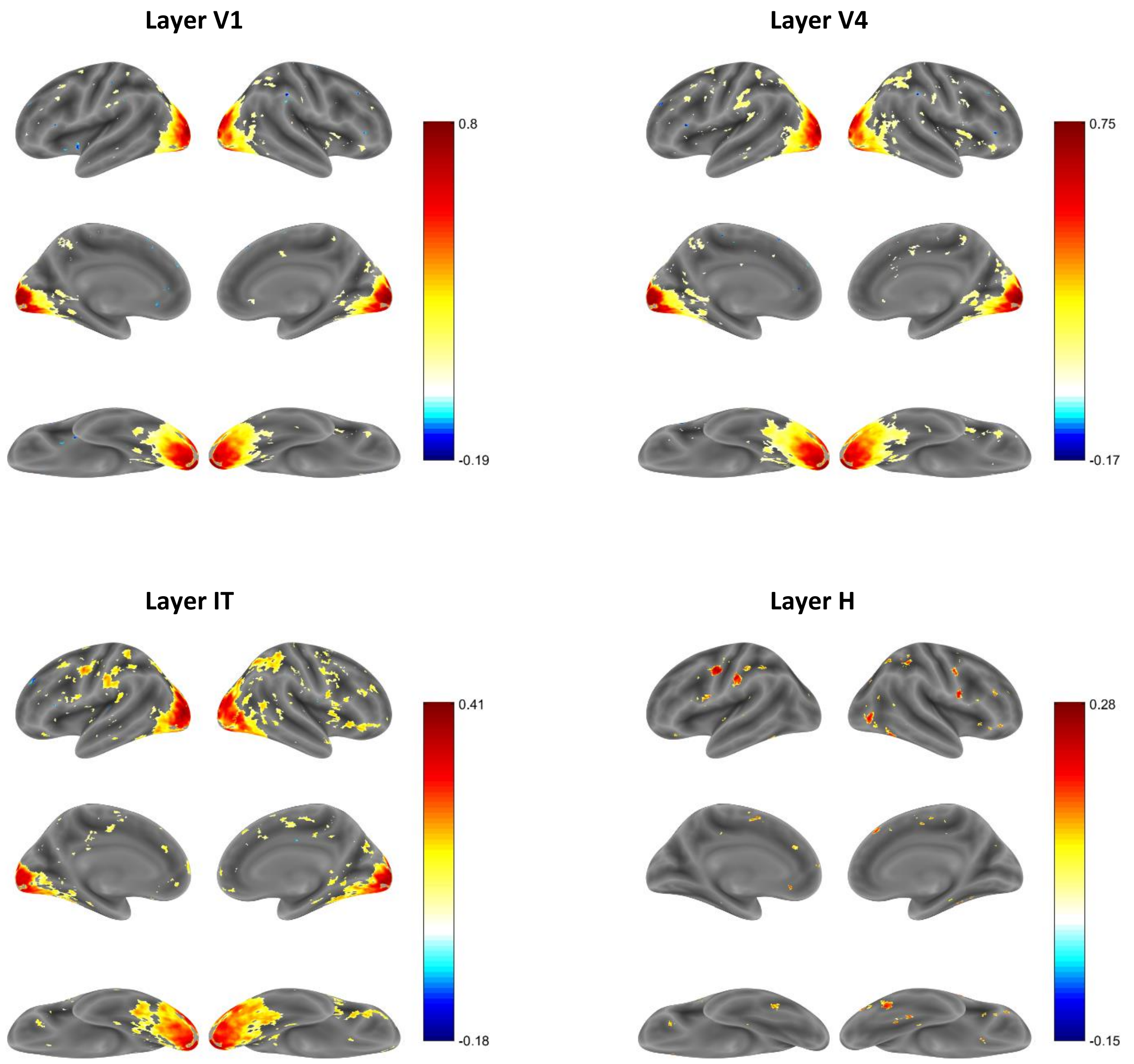


**Figure S12: Searchlight analysis.** Searchlight maps of the correlation between the representation space of CNN layers and neural activity at each voxel. These maps are thresholded to indicate the significant correlation coefficient after FDR correction (p<0.05)

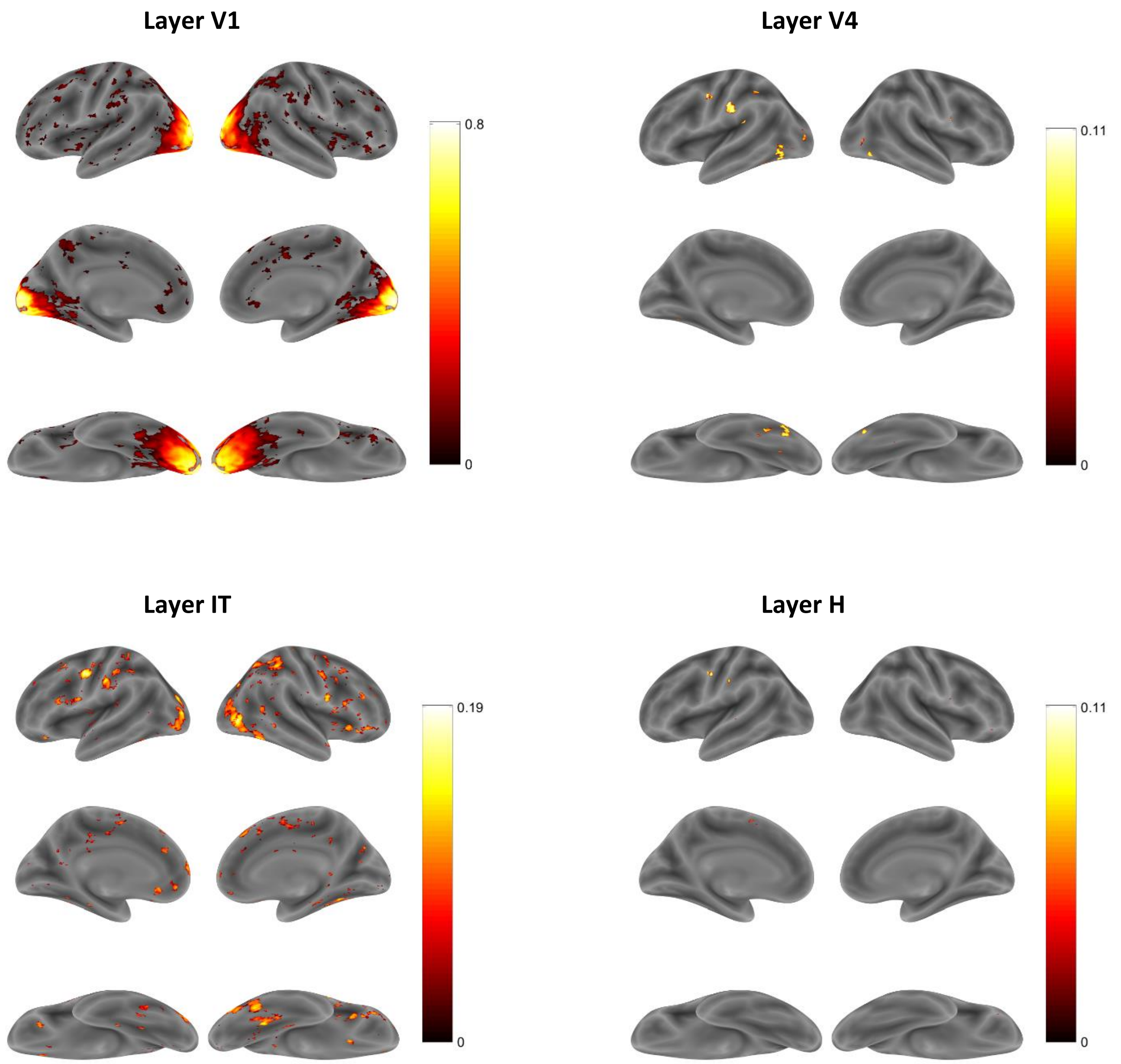


**Figure S13: Searchlight analysis (after step-wise regression).** Searchlight maps of the correlation between the representation space of CNN layers and neural activity at each voxel after step-wise regression. These maps are thresholded to indicate the significant correlation coefficient ($p<0.05$)
